# Enhancement of a STING Agonist Vaccine for Tuberculosis Using Locally Supercharged MS2 Viral Capsids

**DOI:** 10.64898/2026.07.03.736450

**Authors:** Hannah S. Martin, Isabel D. Lamb-Echegaray, Paul Huang, Lillian Shallow, Amir Balakhmet, Preeta Pratakshya, Sarah A. Stanley, Matthew B. Francis

**Author notes:** Corresponding authors. Matthew B. Francis., Sarah A. Stanley. These authors contributed equally to this work (Hannah S. Martin, Isabel D. Lamb-Echegaray).

## Abstract

*Mycobacterium tuberculosis* (Mtb) infection kills more people worldwide than any other pathogen. While the Bacille Calmette-Guérin (BCG) vaccine for Mtb has been widely used for over a century, it provides insufficient protection to eradicate this disease. One of our labs has recently established that a protein antigen (H1) can be combined with a STING pathway agonist to achieve strong protection against Mtb in mice, with performance that exceeds that of the BCG vaccine. However, its reliance on a synthetic cyclic dinucleotide (CDN) with relatively poor cell uptake requires higher dosing levels, thus increasing costs. To increase the efficiency of this vaccine and provide a delivery strategy that could also be used in humans, the H1 Mtb antigen and CDN adjuvant were conjugated to genome-free MS2 viral capsids that included cationic mutations to increase cell uptake. Specifically, the H1 antigen was conjugated to the external surface of MS2 using a tyrosinase-mediated oxidative coupling reaction, and the native STING agonist cGAMP was coupled to internal cysteine residues through a reductively cleavable disulfide linker. The resulting MS2-H1 and MS2-cGAMP conjugates were then co-delivered for three doses of vaccination in mice before exposure to Mtb. The MS2-based vaccine platform was observed to have comparable efficacy to the original H1/CDN formulation, but its enhanced uptake properties enabled 57- to 145-fold less CDN, as well as 3-fold less H1 antigen. Additionally, this vaccine elicited immune responses that have been previously demonstrated to correlate with protection. The ability of the capsid shells to protect the CDN cargo during transport allowed enzymatically produced, and thus readily accessible, cGAMP to be used instead of more costly CDNs that require many synthetic steps. This, combined with the reduced overall amount of CDN that was required, could lower the production costs of future vaccines substantially. Finally, the ability of the capsid-based carriers to bypass the membrane transporters for CDNs suggests that this enhanced vaccination platform is likely to exhibit improved human efficacy in future studies.

## INTRODUCTION

Despite the existence of a vaccine to prevent *Mycobacterium tuberculosis* (Mtb) infection, Mtb continues to be the world’s deadliest single pathogen and has been so for the majority of human history.^1,2^ While this is partially attributed to challenges with antibiotic regimens for treating Mtb infection, a large part of Mtb’s sustained infectious dominance is due the low and inconsistent efficacy of the current Bacille Calmette-Guérin (BCG) vaccine. Unfortunately, BCG has shown between 0 and 80% efficacy in various human vaccination studies.^3,4^ It is moderately effective at preventing severe disseminated tuberculosis disease in children, but there is no strong evidence that it prevents adult pulmonary tuberculosis infections. Taken together, the limitations of the BCG vaccine and the maintained prevalence of Mtb infection point to a strong need for an improved vaccine for this disease.

One of our labs has reported a vaccine for Mtb that outperforms BCG and matches the current top performing Mtb vaccine candidates in mice. For this formulation, a fusion protein antigen for Mtb called H1 was co-delivered with a cyclic-dinucleotide (CDN) stimulator of interferon genes (STING) agonist adjuvant.^5,6^ The STING pathway is an intracellular immune pathway that responds to double-stranded DNA in the cytosol, which makes it particularly relevant for stimulating immune responses against intracellular pathogens like Mtb.^7,8^ The H1/CDN vaccine for Mtb stimulated CD4^+^ T cells and upregulated key immune markers correlated with protection against Mtb: IFN-gamma and IL-17 producing CD4^+^ T-cells. It also increased parenchymal homing CD4^+^ T-cell populations. While highly effective in mice, this vaccine is less likely to show strong efficacy in humans because many CDN-based STING agonists have failed in human clinical trials after showing promising preclinical results.^9,10^ This lack of efficacy in humans is likely at least partially due to inefficient transport of the membrane-impermeable CDN into cells. Additionally, many CDNs are vulnerable to rapid degradation before reaching their targets. We believe that both of these limitations can be addressed by using a viral capsid-based delivery system that can protect internal CDN cargo and increase its cellular uptake.

The bacteriophage MS2 capsid (MS2) is a non-infectious protein-based virus like particle (VLP) that consists of 180 identical coat protein monomers. It is 27 nm in diameter, and it has 32 pores, each 2 nm in diameter.^11,12^ These pores are ideal for the internal loading of small molecule and peptide cargo without requiring a disassembly/reassembly sequence. Additionally, the 2 nm pores are too small for enzymes to enter; thus, MS2 has been demonstrated to protect internal peptide cargo from protease degradation for at least 6 h.^13^ We therefore hypothesized that MS2 could protect internally attached CDN cargo from nuclease degradation in a similar manner, which has recently been shown.^14^

The MS2 capsids used for this vaccine delivery application have been engineered to increase cell uptake. Wild-type MS2 shows poor levels of entry into mammalian cells, but mutants of the capsid with much stronger cell uptake properties have been identified.^13,15,16^ Two point mutations, T71K and G73R (KR), create local positively supercharged regions around the pores (**Figure 1a,b**). Previous evidence supports that these KR mutations likely interact with heparan sulfate motifs on the surfaces of mammalian cells.^13,15,16^ While the mechanism of endosomal escape of MS2-delivered cargo from the endosomes is not fully understood, MS2 capsids with these KR mutations have been shown to deliver multiple types of drug cargo into cells, including drugs with cytosolic targets.^15^ Additionally, these MS2 mutants have been used to deliver peptide antigens, leading to cross presentation in dendritic cells (DC2.4) followed by antigen-specific T-cell activation.^13^ Building upon these previously successful small molecule drug and peptide antigen delivery studies, we report herein the addition of CDN STING agonists as adjuvants, in combination with the H1 fusion protein for efficient Mtb vaccination.

**Figure 1.**
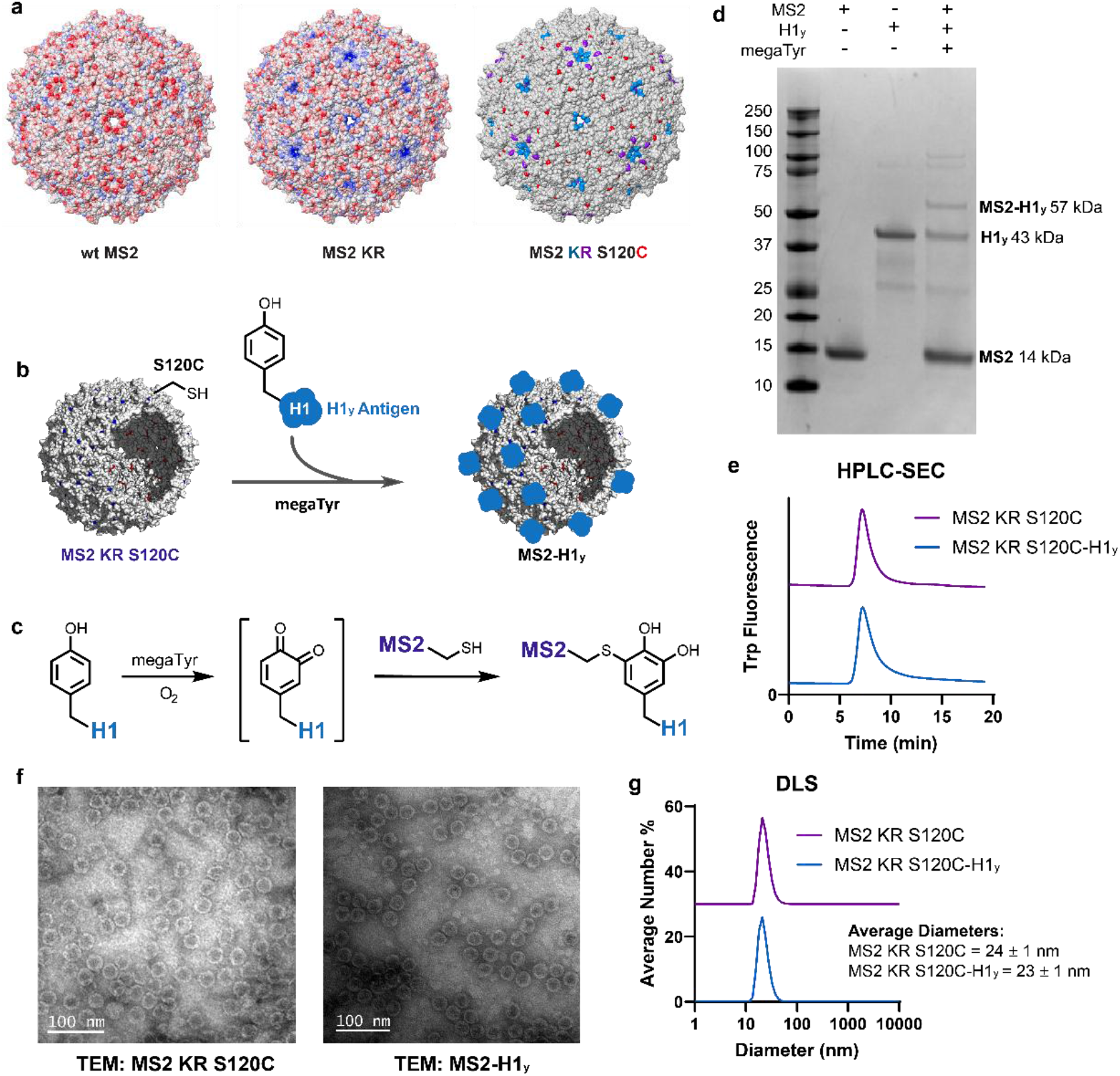
Conjugation of the H1_y_ antigen to MS2 viral capsids for delivery. **(a)** Electrostatic surface potential maps of the wild type bacteriophage MS2 capsid (wt MS2) and MS2 with T71K G73R (MS2 KR) mutations. Positive charges are shown in blue and negative charges are shown in red. Both images use the same scaling for the visualization. A third rendering shows the locations of the external mutations used in this study: T71K is shown in blue, G73R is in purple, and S120C is in red. Capsid images were rendered in Schrödinger/Maestro v 14.4.38. **(b,c)** Schematic of the MS2 + H1y conjugation reaction, which is mediated by the tyrosinase from Bacillus megaterium (megaTyr). **(d)** SDS-PAGE analysis of the MS2 + H1y bioconjugation reaction with expected MWs. **(e)** HPLC-SEC of MS2 KR S120C before and after modification with H1_y_. Assembled capsids elute at ∼7 min. Disassembled capsid proteins would appear after 12 min, but were not observed so long as the level of H1 modification was below 13%. **(f)** TEM images of MS2 KR S120C and MS2 KR S120C-H1y. Scale bars = 100 nm. **(g)** DLS measurements of MS2 KR S120C and MS2 KR S120C-H1y. Average number % is shown.

## RESULTS AND DISCUSSION

The delivery strategy used to build this Mtb vaccine used two different MS2 mutants to allow for external modification of MS2 with H1 and internal modification of MS2 with the cyclic guanosine monophosphate–adenosine monophosphate (cGAMP) STING agonist adjuvant. For H1 conjugation to MS2, the KR S120C MS2 mutant was used, (**Figure 1a,b**). As previously described, the KR mutations are believed to bind heparan sulfate on cells and allow for potent cell uptake of MS2. The additional S120C mutation is an external cysteine that was used for external conjugation of H1. For adjuvant delivery, the cGAMP STING agonist required internal conjugation to MS2 to protect it from nuclease degradation. Following our previously reported delivery of internally conjugated peptide antigens in MS2, the S37P KR MS2 (PKR MS2) mutant was used for cGAMP delivery.^17^ This mutant also contains an internal N87C mutation for site-specific internal modification (**Figure 2a,e**).

**Figure 2.**
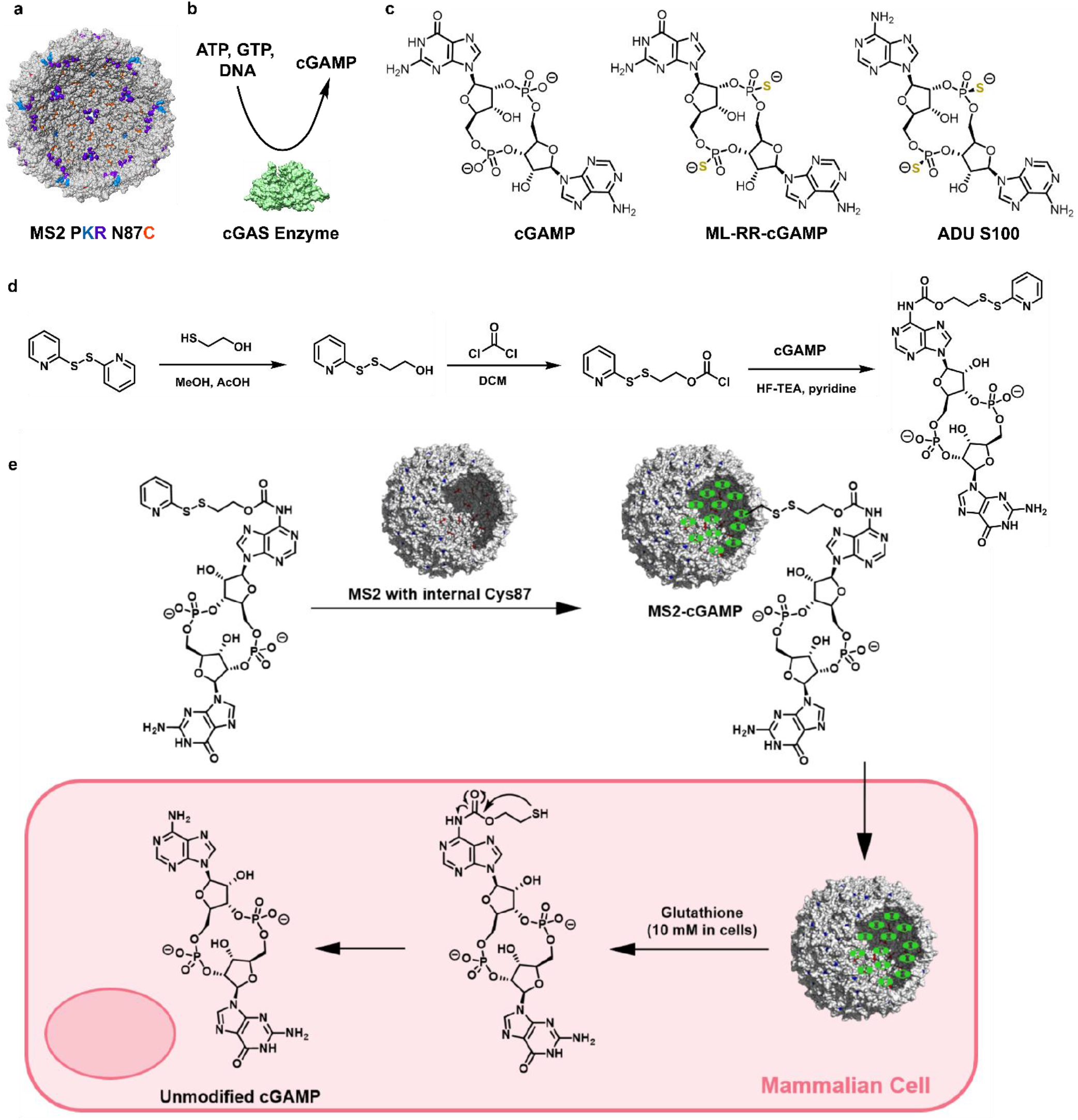
Design and synthesis of cGAMP-loaded PKR MS2 capsids. **(a)** Rendering of the internal surface of MS2 PKR N87C capsids, with the N87C mutations shown in orange, T71K mutations in blue, and G73R in purple. The stmcture was generated in Schriidinger/Maestro v 14.4.38. **(b)** Structures of the native STING agonist, cGAMP, and a synthetic STING agonist drug, ML-RR-cGAMP. **(c)** Enzymatic synthesis of cGAMP was achieved using recombinantly expressed cGAS enzyme. **(d)** Synthesis of a disulfide linker and its attachment to cGAMP. **(e)** Addition of the cGAMP-linker construct to MS2 and mechanism of traceless cGAMP release after reduction by glutathione in cells.

### Conjugation of the H1 Antigen to MS2 Capsids

Expanding on our previous results, which have shown successful adaptive immune activation in response to MS2- delivered peptide antigens in cell culture,^17^ we first conjugated the H1 Mtb antigen to the external surface of MS2 capsids. The H1 antigen is a 43 kDa fusion protein of Ag85b and ESAT-6, two immunodominant proteins for Mtb.^18,19^ Ag85b is the most abundant periplasmic Mtb protein and is essential for cell envelope synthesis.^20^ ESAT-6 is a virulence factor that is abundantly secreted by Mtb. During an active Mtb infection, Ag85b-specific CD4^+^ T-cell populations peak early after infection and decrease over time, while the ESAT-6-specific CD4^+^ T-cell population rises in the beginning of an infection and maintains high levels throughout its course.^21^ Therefore, both antigens play an important role in the immune response against Mtb. The H1 antigen is well established to provide protection against tuberculosis infection in many animal and non-human primate models.^22–27^

To allow for bioconjugation with MS2, the H1 fusion protein antigen was recombinantly expressed in *Escherichia coli* (*E. coli*) with a C-terminal (SG_4_)_4_Y sequence (H1_y_). This provided a flexible extension followed by a tyrosine residue that was adequately exposed for a tyrosinase-based bioconjugation approach.^28^ While following a previously established soluble protein purification method yielded pure H1_y_, the yield obtained was low (∼0.5 mg/L of *E*.*coli* culture).^23^ The majority of H1_y_ protein was trafficked to inclusion bodies of *E. coli*, so a new insoluble protein purification method was developed to increase the antigen yield. This method involved washing and then solubilizing the inclusion bodies, followed by a denaturation/refolding sequence. Cleavage of the His_6_ purification tag using TEV protease generated H1_y_ with a 100× increase in protein yield (∼50 mg/L of *E. coli* culture) compared to the previously established soluble purification.

The H1_y_ antigen was covalently attached to the external surface of KR S120C MS2 for delivery into cells using an enzymatic oxidative coupling reaction.^28^ This reaction used a recombinantly expressed tyrosinase enzyme from *Bacillus megaterium* (megaTyr), which selectively oxidizes only highly surface exposed tyrosine residues.^29^ The tyrosinase oxidative coupling reaction proceeded through two steps: first, the terminal tyrosine was oxidized to form an *ortho*-quinone intermediate and second, the cysteine nucleophile on MS2 added to form the catechol product in an enzyme-independent step (**Figure 1b, c**). The resulting aryl-thiol linkage formed by tyrosinase couplings has proven to be irreversible in serum.^28^ However, once inside cells it is believed that proteases release H1_y_ from MS2 similarly to previously demonstrated peptide antigen release from MS2.^13^ This conjugation method is particularly attractive because it utilizes two natural amino acids rather than requiring non-canonical amino acid incorporation like many other site-specific bioconjugation methods. Additionally, the megaTyr enzyme only activates terminal and highly surface-exposed tyrosine residues, so it does not interfere with the native tyrosines in most proteins (including H1_y_).

After performing the KR S120C MS2-H1_y_ (MS2-H1_y_) conjugation, it was vital that the megaTyr enzyme was completely removed from the product because any residual oxidation was observed to cause toxicity in cell culture. To ensure complete removal of the enzyme, MS2-H1_y_ conjugation reactions were performed using a His_6_-tagged megaTyr enzyme that was immobilized on nickel resin. When the reactions were complete, they were filtered to remove the resin-bound enzyme as confirmed using previously reported techniques.^13^ It is also important to note that the MS2-H1_y_ tyrosinase coupling reactions were performed at pH 8 to ensure that megaTyr maintained strong binding to the resin because the strength of the His_6_-nickel affinity interaction is pH dependent.^30^ Lower buffer pH values, such as pH 6.5, have led to partial release of megaTyr from the resin.

SDS-PAGE analysis using a reducing loading buffer that contained 10 mM DTT confirmed that the MS2-H1_y_ linkage was the expected oxidative coupling product rather than a disulfide exchange product with a cysteine on H1_y_ (**Figure 1d**). Because the H1_y_ antigen (∼43 kDa) is much larger than each of the 180 MS2 monomers in the capsid (∼14 kDa), there was a maximum threshold of H1_y_ loading before the MS2 capsid was destabilized by steric repulsion of the antigens. The limit to H1_y_ loading on MS2 before the capsid disassembly was approximately 25 copies of H1_y_ per capsid (13% H1_y_ on MS2), as estimated by densitometry of the SDS-PAGE and HPLCSEC confirmation of capsid assembly state (**SI Figure S2**). Additionally, SEC-HPLC was used to analyze if the MS2-H1_y_ conjugates stayed assembled over time when stored at 4 °C, revealing no signs of disassembly after more than three weeks (**SI Figure S3**). PEG linkers between H1_y_ and MS2 were explored to test if longer linkers would release some steric strain and increase the amount of H1_y_ that could be loaded on MS2, but the linkers did not improve the loading efficiency (**SI Figure S4**). A specific stoichiometric ratio of MS2 to H1_y_ was used to control the reaction and prevent conjugation exceeding 13% (∼25 copies per capsid) for all MS2-H1_y_ conjugates used in vaccination studies.

HPLC-SEC, TEM, and DLS were used to ensure MS2-H1_y_ conjugates maintained the capsid assembly state. Assembled MS2 capsids elute from HPLC-SEC around 7 min, while disassembled MS2 oligomers and monomers elute around 12 and 16-18 min, respectively. Disassembly products were not observed (**Figure 1e**). Additionally, fully assembled capsids were observed by DLS and TEM, with average diameters within the expected ranges for MS2 capsids (**Figure 1f, 1g**). Because the level of H1_y_ labeling on MS2 capsids was kept below ∼13%, little difference between the MS2 vs. MS2-H1_y_ capsid diameters was observed by TEM and DLS.

### Synthesis and Delivery of MS2-cGAMP for STING Activation

It is well appreciated that vaccines require adjuvants to stimulate the immune response to an antigen, so the second component of the vaccine focused on adjuvant delivery. While one of our labs has shown that using 5 µg of a STING agonist CDN called ML-RR-cGAMP (2′–5′ and 3′–5′ (mixed linkage)- RR-cyclic guanosine monophosphate-adenosine monophosphate with phosphorothioate linkages for phosphatase resistance, structure in **Figure 2c**) is effective as an adjuvant for Mtb vaccination in mice,^5,6^ there are some key limitations to this approach. First, the relatively high dose of this small molecule is likely required because the drug is cell membrane-impermeable and must rely on low-affinity cell-surface anion transporters to enter relevant immune cells.^31–33^ This effect is magnified as the drug is degraded outside the cell *in vivo*, albeit somewhat more slowly due to its phosphorothioate linkages as phosphate replacements. Second, CDNs including ML-RR-cGAMP are expensive, rendering high doses impractical for worldwide vaccine distribution. With these limitations in mind, we aimed to use MS2 to deliver a much lower dose of STING agonist adjuvant and bypass the need for CDN transporters.

We chose the native STING agonist cGAMP as our adjuvant because it could be synthesized enzymatically using recombinantly expressed cyclic GMP-AMP synthase (cGAS) enzyme (**Figure 2b,c**).^34,35^ To do this, recombinantly expressed cGAS enzyme was mixed with ATP, GTP, and DNA. The cGAS-synthesized cGAMP was then purified from the reaction for vaccine incorporation using reversed phase chromatography.

The next step was the covalent attachment of cGAMP to the internal surface of MS2 for delivery. A self-immolative disulfide linker was used as previously.^14,16^ After chemical synthesis of the linker, the chloroformate group was reacted with the adenosine amino group of cGAMP to form a carbamate product (**Figure 2d**). This cGAMP-linker conjugate was designed to disulfide exchange with cysteine residues inside PKR MS2. Once delivered to cells, the disulfide bond between PKR MS2 and cGAMP was envisioned to be cleaved by glutathione in the cell cytoplasm.^16^ Once cleaved, the linker was designed to self-immolate and release cGAMP with no residual “scar” (**Figure 2e**).^14^ To our knowledge this is the only delivery strategy that can release cGAMP from a covalent linkage without structural alteration.

The cGAMP-linker conjugate was subjected to disulfide exchange with an internal C87 mutation in PKR MS2 (**Figure 2a,e**). The product of this reaction was analyzed using LC-MS (ESI-TOF), which indicated that PKR MS2 was highly labeled with cGAMP (**Figure 3a**). This reaction reliably yielded between 60% and 85% conjugation of cGAMP on PKR MS2, corresponding to 108 to 153 molecules of cGAMP per capsid. To ensure MS2 maintained its assembled state after conjugation, HPLC-SEC was used to analyze the product (**Figure 3b**). The resulting chromatogram showed only one population with an elution time around 7 min, corresponding to assembled capsids. Additionally, TEM and DLS measurements indicated spherical capsids with diameters in the expected range (**Figure 3c,d**).

**Figure 3.**
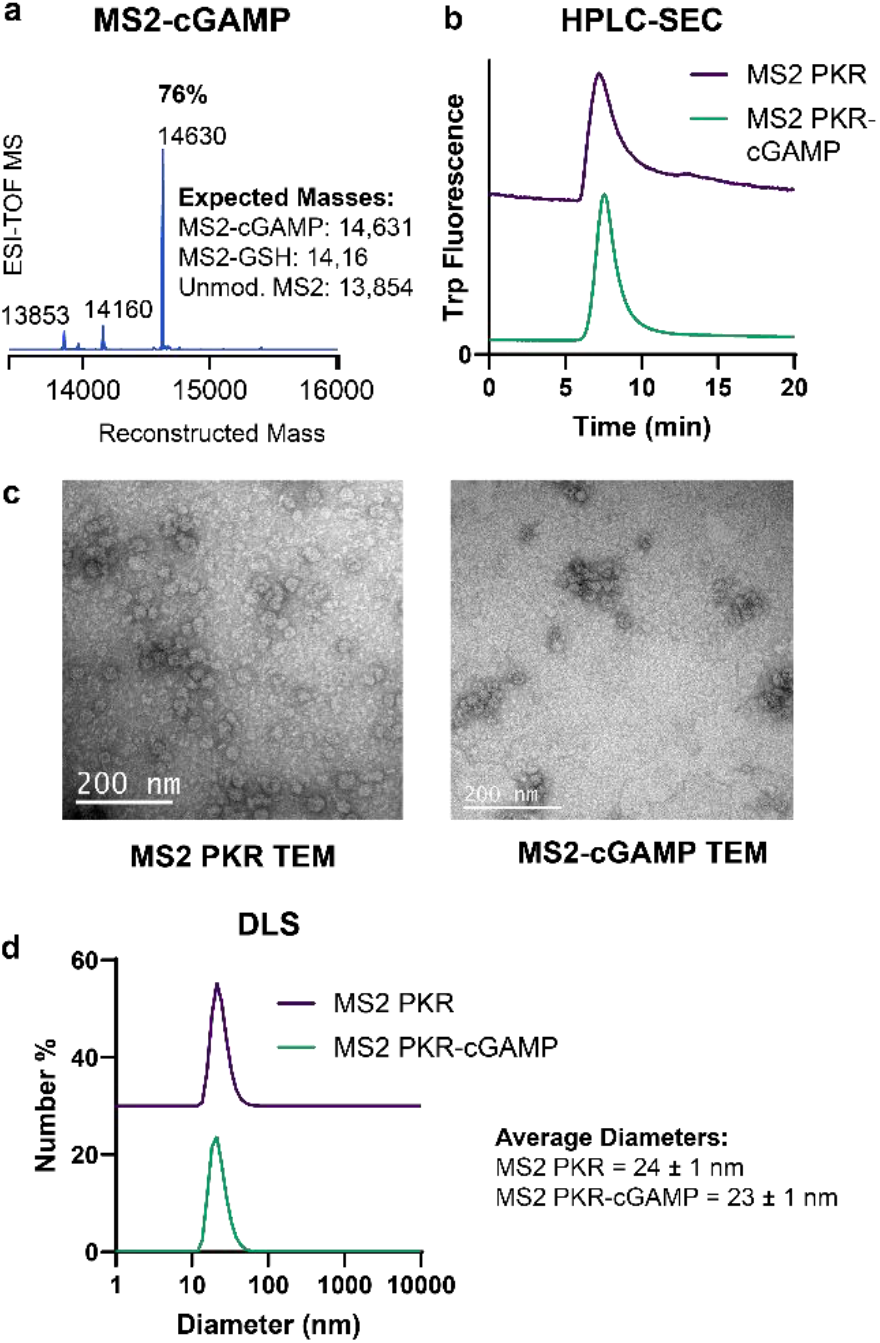
Characterization of MS2 PKR-cGAMP vehicles. **(a)** LC ESI-TOF MS analysis of the MS2 PKR-cGAMP product.**(b)** HPLC-SEC analysis of assembled MS2 PKR and MS2 PKR-cGAMP capsids. Disassembled capsids would appear after 12 min, but were not observed.**(c)** TEM images of MS2 PKR and MS2 PKR-cGAMP. Scale bars = 200 nm.**(d)** DLS measurements of MS2 PKR and MS2 PKR-cGAMP capsids. Average number % is shown.

Previously reported *in vitro* STING activation using MS2-cGAMP compared to non-delivered native cGAMP and a pharmaceutical CDN STING agonist with phosphorothioate linkages that is very similar to ML-RR-cGAMP (ADU S100, **Figure 2c**) support the idea that a substantially lower dose of STING agonist could be used to vaccinate against Mtb with MS2 delivery.^14^ Using a THP-1 monocyte STING reporter cell line for a dose response measurement, MS2-cGAMP was compared to free cGAMP and free ADU S100 for STING activation. The EC_50_ value measured for MS2-cGAMP was approximately 22 nM, while ADU S100 was about 9960 nM, more than 450-fold higher despite the nuclease-resistant phosphorothioate linkages that ADU S100 contains. Due to degradation by serum proteins, the cGAMP response was exceptionally poor, exceeding 50,000 nM. Therefore, MS2- cGAMP delivery led to an over 2,000-fold increase in the cGAMP drug’s STING activation potency.^14^ These data suggest that with MS2 delivery, magnitudes less expensive STING agonist adjuvants could be used to produce the same vaccine adjuvant immune response. This quality is particularly valuable when cost per dose of the vaccine is a key factor in allowing for global distribution. Further, parallel studies using CDN transporter (SLC19A1) knockout cell lines have shown that MS2 delivery does not require these CDN transporters for the cGAMP to enter cells and activate the STING pathway.^14^ This is particularly relevant because different cell types and species present varying amounts of cell surface anion transporters and have different CDN uptake mechanisms, which could lead to inconsistent STING activation from CDN drugs that are reliant on transporters.^31–33^ Because MS2-cGAMP delivery does not rely on transporters, this strategy of delivering the vaccine adjuvant is significantly more likely to translate to comparable immune activation in humans.

### MS2 Delivery of H1/cGAMP Controlled Infection Upon *M. tuberculosis* Challenge at Low Vaccine Concentrations

In previous studies we showed that a high dose of the cyclic dinucleotide (CDN) ML-RR-cGAMP, combined with a fusion protein antigen composed of two highly immunogenic tuberculosis antigens Ag85b and ESAT6 (H1), provided strongly effective protection against tuberculosis in mice when administered in a three-dose regimen.^5,6^ Hereafter, this treatment is referred to as H1/CDN. However, despite the increased serum stability of ML-RR-cGAMP compared to native cGAMP, protection was achieved at relatively high H1 and cGAMP concentrations.

Therefore, we tested whether using the MS2 viral capsid to deliver the antigen and adjuvant (MS2-H1_y_/MS2- cGAMP) could enhance vaccine potency and enable protection against Mtb at reduced doses of each vaccine component. Mice were immunized intranasally (i.n.) with PBS as a negative control, the previously established dose of H1/CDN as a positive control, the MS2-H1_y_/MS2-cGAMP vaccine, an equivalent concentration of H1/cGAMP without MS2 delivery, or MS2-H1_y_ alone. Vaccinations were administered in a three-dose regimen at four-week intervals. Four weeks after the final immunization, the mice were challenged with *M. tuberculosis* strain Erdman, and protective efficacy was assessed by measuring lung bacterial burden by enumerating colony forming units (CFU) four weeks post-challenge **(Figure 4a)**.

**Figure 4.**
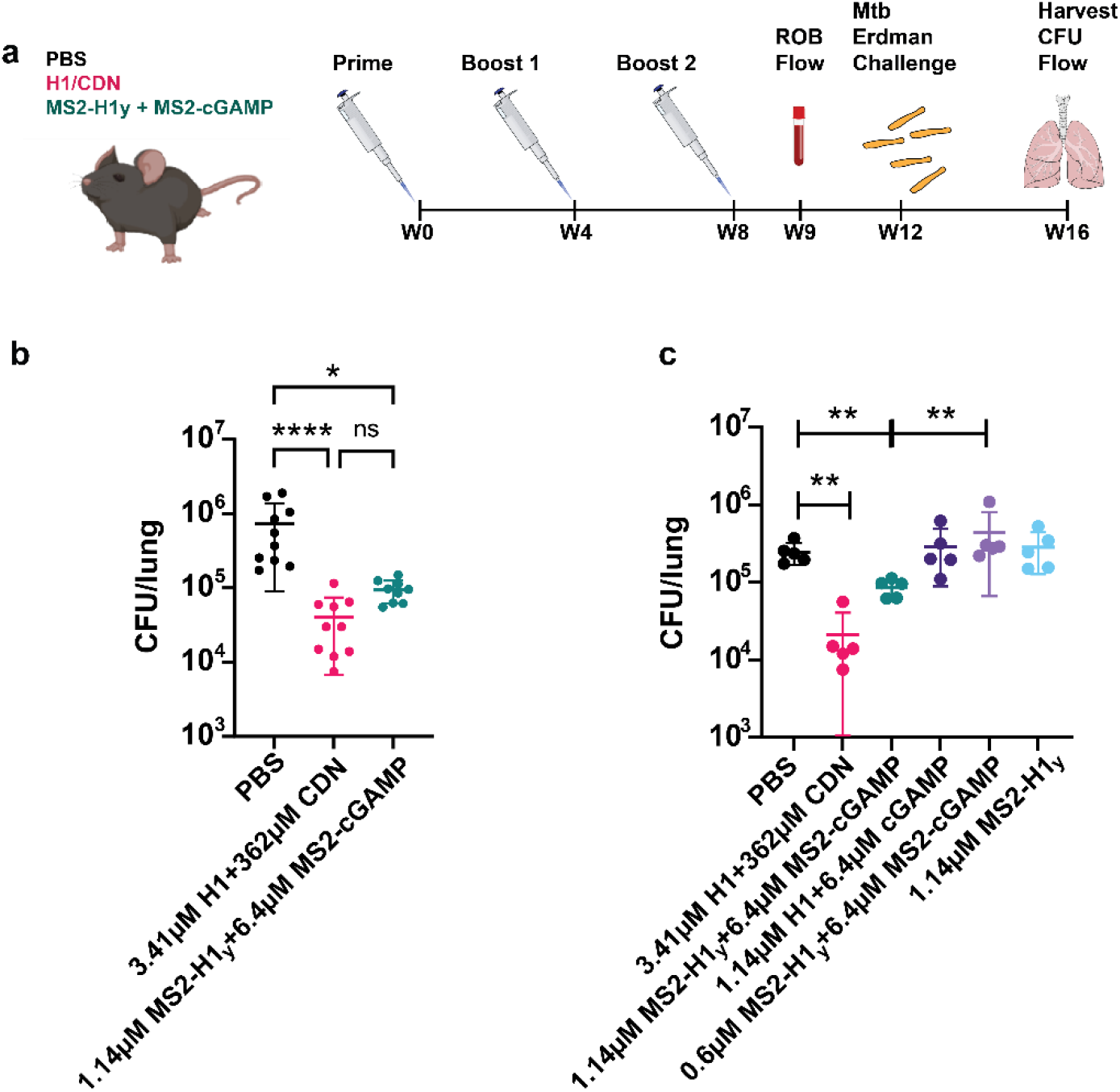
Vaccination with MS2-H1y + MS2-cGAMP by an i.n. route elicits protection at 57x lower concentrations of CDN and 3x lower concentrations of H1. (a) Experimental procedure and vaccine schedule. Mice were vaccinated intranasally three times at 4-wk intervals with either H1/CDN or MS2-H1y + MS2-cGAMP 12 weeks before challenge. At 4 weeks post challenge, the mouse lungs were harvested and analyzed for bacterial burden. (b) CFUs were measured for mice vaccinated with either H1/CDN or MS2-H1y + MS2-cGAMP at 4 weeks post challenge, compared to a PBS vehicle. Data are expressed as mean (±SD) of eight to ten mice per group from two pooled, independent experiments. The values corresponding the individual animals are also displayed. Kruskal-wallis test p values; *p ≤ 0.05, **p ≤ 0.01, ***p ≤ 0.001. (c) CFUs are summarized for control groups. Mice vaccinated with either high or low concentrations of free H1/CDN, low concentrations of MS2-H1y + MS2-cGAMP, or only MS2-H1 were evaluated 4 weeks post challenge. Data are expressed as mean (±SD) of five mice per group, with the values from individual animals displayed. *p ≤ 0.05, **p ≤ 0.01, Mann-Whitney t test. Panel a created with BioRender.com.

Immunization with MS2-H1_y_/MS2-cGAMP conferred robust protection, resulting in ∼1 log lower CFU/lung than the PBS control, at 57-fold and 145-fold lower concentrations of CDN adjuvant and a 3-fold lower concentration of H1 antigen when compared to the previously established H1/CDN vaccine. Despite the substantially lower doses of MS2-delivered antigen and adjuvant, the protection against Mtb infection was approximately equivalent to the protection elicited by H1/CDN **(Figure 4b, SI Figure S6)**. Notably, this was achieved with native cGAMP, which is more cost effective to produce but lacks the phosphorothioate linkages of ML-RR-cGAMP that provide serum stability. As MS2 both shields native cGAMP from degradation and enhances its delivery to relevant immune cells, it allows for the effective usage of native cGAMP at such a low concentration. This was demonstrated through immunization with the equivalent lower concentrations of H1 and cGAMP without MS2 delivery. This condition failed to protect against Mtb challenge, demonstrating the MS2 delivery platform is essential to achieve lower effective dosages of vaccine and adjuvant (**Figure 4c, SI Figure S6**).

To determine minimally sufficient components of the vaccine formulation, MS2-H1_y_ or MS2-cGAMP were administered independently and failed to protect against Mtb challenge, demonstrating that improved delivery of either H1 or cGAMP alone is insufficient for protection and that both components are required in combination. Additionally, a lower concentration of MS2-H1_y_ (0.6µM) with MS2-cGAMP also failed to provide protection, suggesting that the minimum protective concentration of H1 lies above 0.6 µM (**Figure 4c, SI Figure S6**).

These data demonstrate that MS2 enhanced vaccine delivery, substantially reducing the concentrations of both antigen and adjuvant required for protection. Notably, protection was achieved without nuclease-resistant phosphorothioate linkages on cGAMP, likely because MS2 protected cGAMP from *in vivo* enzymatic degradation. MS2- cGAMP delivery also likely increased the rate of cell uptake of cGAMP because the engineered MS2 mutants have their own cell-uptake properties that bypass the need low-affinity and inconsistently expressed cell-surface for CDN transporters. These findings highlight the effectiveness of a MS2-delivered Mtb protein subunit vaccine. The bioconjugation chemistry and viral capsid delivery system presented herein enabled the use of remarkably low doses of antigen and adjuvant with potential for human efficacy, suggesting the possibility of affordable global distribution.

### MS2 Delivery of H1 and cGAMP Elicited Protective IL-17 and IFN-γ CD4 T cells at Low Vaccine Concentrations

CD4^+^ (helper) T cells are essential to the protective immune response against tuberculosis. We showed previously that protection elicited by the H1/CDN vaccine requires both IL-17– and IFN-γ–cytokine producing CD4^+^ T cells, and also that vaccination generates protective populations of each in the blood, with influx and expansion of both subsets in the lungs upon Mtb challenge.^5,6^ Therefore, we analyzed levels of these cytokine-producing T cells in MS2-H1_y_/MS2-cGAMP vaccinated mice. We showed that the H1/CDN vaccine elicited protective IL-17 and IFN-γ producing CD4^+^ T cells in the blood of vaccinated mice and caused influx and expansion of these T cell populations in the lungs upon Mtb challenge. To determine whether MS2 delivery of H1 and cGAMP produced similar cytokine responses to the H1/CDN vaccine, even with much lower antigen and adjuvant concentrations, we measured the numbers of Ag85B and ESAT6-specific (each component of the H1 fusion protein antigen) IL-17 and IFN-γ producing CD4^+^ T cells in the blood (**Figure 5**) and lungs (**Figure 6**) of vaccinated mice. The blood samples, collected 3 weeks prior to Mtb challenge, and the lung samples, collected 4 weeks post-Mtb challenge (**Figure 4a**), were analyzed using intracellular cytokine staining (ICS) and flow cytometry.

**Figure 5.**
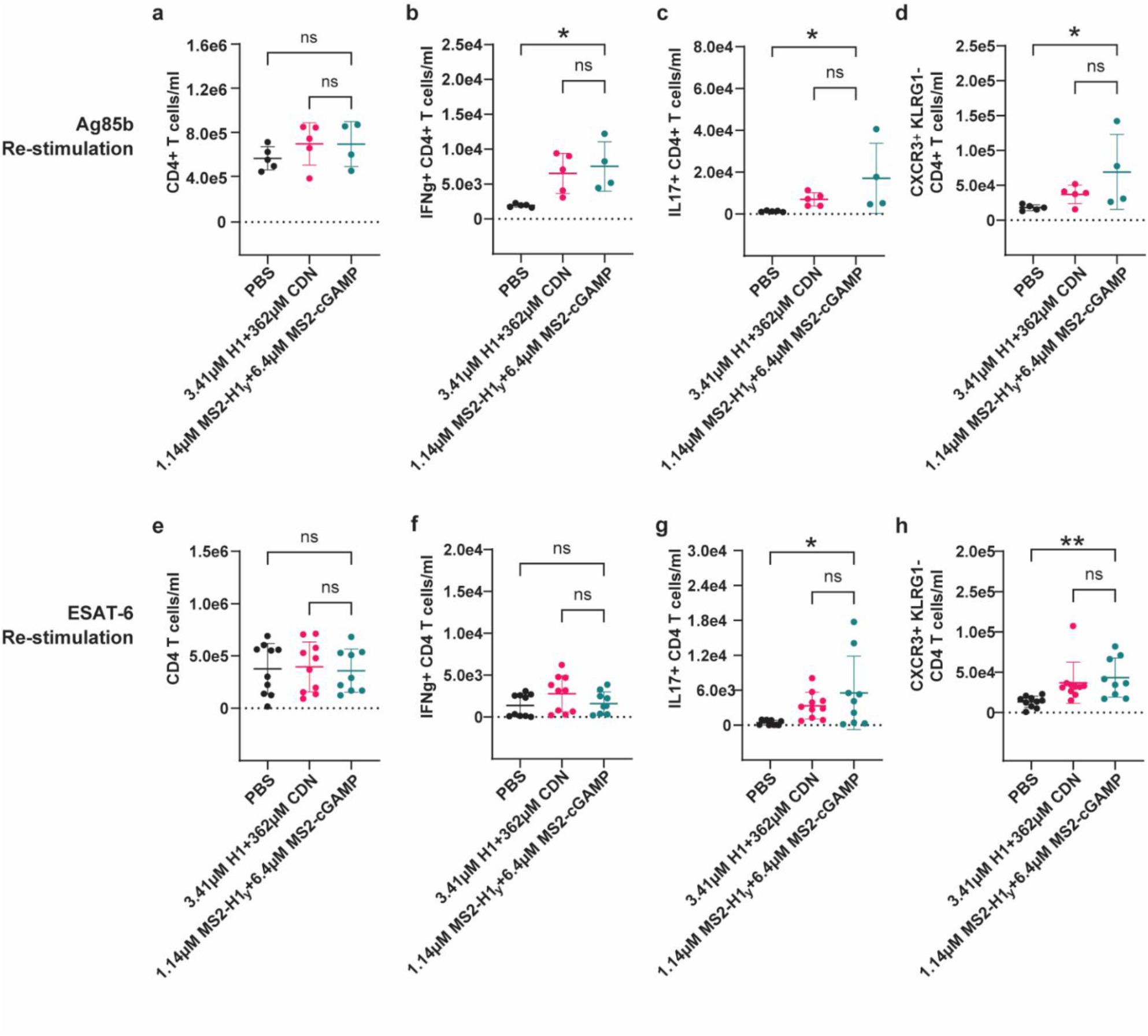
Flow cytometry analysis of immune responses from blood taken from mice vaccinated with HI/CD or low concentrations ofMS2-Hly/MS2-cGAMP. Wild-type C57BL/6 (B6) mice were vaccinated intranasally three times at 4 week intervals with either H 1/CDN or MS2-H ly/MS2-cGAMP. Three weeks prior to challenge, blood was collected and processed before and after re-stimulation with either Ag85b or ESAT-6. Flow cytometry data are shown for CD4 T cell concentrations after re-stimulation with (a) Ag85b or (e) ESAT-6. !CS of blood CD4 T cells that produce IFNg after re-stimulation with (b) Ag85b or (f) ESAT-6. JCS of CD4 T cell counts that produce JL17 after re-stimulation with (c) Ag85b or (g) ESAT-6. CD4 T cell counts that are CXCR3+ and KLRG I-after re-stimulation with (d) Ag85b or (h) ESAT-6. Data are expressed as mean (± SD) of 4 to ten mice per group from one representative experiment or two independent, pooled experiments. The values corresponding to individual animal are di played. Kruskal-Wallis test p values; *p ≤ 0.05, **p ≤ 0.01, ***p ≤ 0.001.

**Figure 6.**
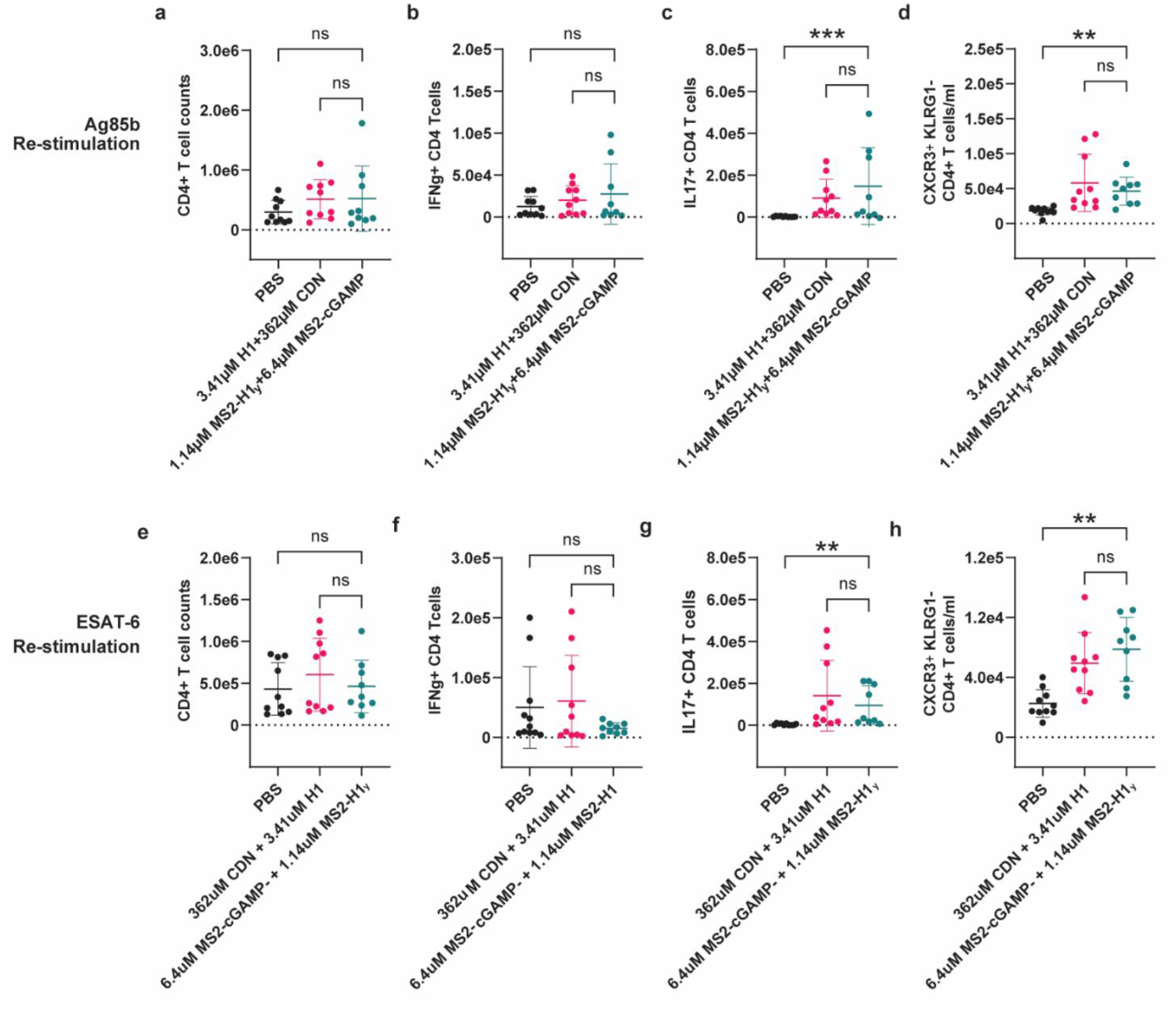
Flow cytometry analysis of immune responses in the lungs ofmice vaccinated with Hl/CD or low concentrations ofMS2-H Ly+ MS2-cGAMP. Wild-type C57BL/6 (B6) mice were vaccinated intranasally three times at 4 week .intervals with either Hl/CDN or MS2-Hly/MS2-cGAMP. At 4 weeks post challenge mouse lungs were harvested, processed for flow cytometry and re-stimulated with either Ag85b or ESAT-6. Flow cytometry data are shown for CD4 T cell counts after (a) Ag85B re-stimulation or (e) ESAT-6 re-stimulation, lCS of lung CD4 T cells that produce IFNg after (b) Ag85B re-stimulation or (f) ESAT-6 re-stimulation, ICS of CD4 T cell counts that produce IL 17 after (c) Ag85B re-stimulation or (g) ESAT-6 re-stimulation, and CD4 T cell counts that are CXCR3+ and KLRG 1- after (d) Ag85B re-stimulation or (h) ESAT-6 re-stimulation. Data are expressed as mean (± SD) of eight to ten mice per group from two independent, pooled experiments with values from individual animals displayed. Kruskal-Wallis test p values; *p ≤ 0.05, **p ≤ 0.01, ***p ≤ 0.001.

The measured immune responses in the blood and lungs were very similar between the H1/CDN vaccine and the MS2-H1_y_/MS2-cGAMP vaccine. At 3 weeks prior to Mtb challenge, we found that there were no statistical differences in the numbers of CD4^+^ T cells in the blood of H1/CDN, MS2- H1_y_/MS2-cGAMP, or PBS vaccinated mice (**Figure 5a, 5e**). However, both H1/CDN and MS2-H1_y_/MS2-cGAMP vaccines elicited higher concentrations of Ag85b-specific IFN-γ producing CD4^+^ T cells in the blood relative to PBS (**Figure 5b**). Interestingly, some groups did not have higher ESAT-6- specific IFN-γ producing CD4^+^ T cells (**Figure 5f**). This was the only immune response in the blood or lungs of vaccinated mice that was observed to favor either the Ag85b or ESAT-6 antigen specifically (**Figure 5, 6**). At the peak of the immune response, 4 weeks after challenge, there were no significant differences in either the number of CD4^+^ T cells (**Figure 6a, 6e**) or the IFN-γ-producing CD4^+^ T cells (**Figure 6b, 6f**) in the lungs of mice vaccinated with either H1/CDN, MS2-H1_y_/MS2- cGAMP or PBS. However, both H1/CDN and MS2-H1_y_/MS2- cGAMP vaccines elicited significantly higher numbers of IL-17-producing CD4^+^ T cells in the blood (**Figure 5c, 5g**) of vaccinated mice relative to PBS prior to Mtb challenge. These cells were especially elevated in the lungs (**Figure 6c, 6g**) after Mtb challenge, suggesting that in these vaccinated mice, relevant immune cell populations migrated to the lungs in response to the Mtb challenge. Importantly, the lower concentration doses of the MS2-H1_y_/MS2-cGAMP vaccine produced essentially identical Mtb-protective immune responses as the higher concentration H1/CDN vaccine, with no statistically significant differences in the numbers of CD4^+^ T cells and IFN-γ or IL17 producing CD4^+^ T cells in the blood or lungs of vaccinated mice.

These data demonstrate that MS2-delivery of H1 and cGAMP elicited protective IL17 and IFN-γ producing CD4^+^ T cells in blood after vaccination and influx and expansion of these immune responses in the lungs after Mtb challenge. In addition, the immune responses produced by MS2 delivery of H1 and cGAMP were strikingly similar to those elicited by the previously established H1/CDN vaccine, despite substantially lower concentrations of both antigen and adjuvant in MS2. Further, the MS2-delivered vaccine provided equal immune responses to the previously established vaccine while using the much more cost-effective cGAMP adjuvant, which does not have any nuclease resistance properties, because the MS2 capsid is nuclease resistant.

### MS2 delivery of H1 and cGAMP Increased Parenchyma Homing T Cells Relative to PBS Vaccinated Mice

It has previously been demonstrated that CD4^+^ T cells localized to the lung tissues (CD4^+^CXCR3^+^KLRG1^−^) are correlated to protection against Mtb in mice. These cells have been described to home to the lung parenchyma and are protective against Mtb infection when adoptively transferred into mice.^36^ We have previously shown that vaccination with H1/CDN resulted in an increase in these CD4^+^CXCR3^+^KLRG1^−^ T cells.^5,6^ Therefore, we tested if MS2- delivery of H1 and cGAMP elicited similar immune responses even with lower antigen and adjuvant concentrations than H1/CDN. The numbers of Ag85b and ESAT6-specific CXCR3^+^ KLRG1^−^ CD4^+^ T cells in the blood and lungs of vaccinated mice were measured before and after Mtb challenge by flow cytometry.

At 3 weeks pre-challenge, we observed a higher concentration of CD4^+^CXCR3^+^KLRG1^−^ T cells in the blood of mice vaccinated with either H1/CDN or H1-MS2/cGAMPMS2 compared to PBS (**Figure 5d, 5h**). At 4 weeks after challenge, both H1/CDN and MS2-H1_y_/MS2-cGAMP vaccinated mice also had increased lung parenchyma homing CD4^+^CXCR3^+^KLRG1^−^ T cells compared to the PBS control group (**Figure 6d, 6h**). These immune responses elicited by H1/CDN and MS2-H1_y_/MS2-cGAMP were highly comparable, with no statistically significant differences in the numbers of CD4^+^CXCR3^+^KLRG1^−^ T cells in the blood or lungs of the vaccinated mice. Additionally, the immune responses to restimulation with Ag85B or ESAT-6 were similar, demonstrating that both components of the H1 antigen were involved, and neither specific antigen dominated the response (**Figure 5d, 5h, 6d, 6h**).

Taken together, these data demonstrate that MS2- delivery of H1 and cGAMP elicited expansion of protective lung parenchyma homing CD4^+^ T cells in the blood after vaccination, and these T cell populations indeed exhibited elevated levels in the lungs after challenge with Mtb. The immune responses produced by MS2-delivered H1 and cGAMP were equivalent to those elicited by H1/CDN, even with substantially lower concentrations of both antigen and adjuvant and in the absence of nuclease resistant phosphorothioate linkages on cGAMP.

## CONCLUSION

In this study, we have improved upon an effective, previously reported Mtb vaccine using a viral capsid-based delivery system. This new platform achieved comparable levels of protection and immune activation in mice to those of our previously reported combination of the H1 antigen and MLRR-cGAMP STING agonist adjuvant. Notably, however, the enhanced cell uptake of the capsid mutants increased the dosing efficiency substantially. Notably, however, the enhanced cell uptake of the capsid mutants increased the dosing efficiency substantially. The MS2-H1_y_/MS2-cGAMP system achieved a 57- to 145-fold reduction in the amount of CDN adjuvant that was required, as well as a 3-fold reduction in the amount of H1 antigen. Moreover, the ability of the capsid shells to protect the CDN cargo during transport allowed for enzymatically produced, and thus readily accessible, cGAMP to be used instead of more costly CDNs that require many synthetic steps. Additionally, the MS2 capsid-based carriers for CDNs bypass membrane transporters, leading to more efficient and consistent adjuvant delivery that is likely to translate to human efficacy. These benefits could lower the production costs of future vaccines substantially.

Several of the immunological signaling outcomes observed in this study reached statistical significance, demonstrating the capacity of the vaccine platform to elicit robust immune responses. At the same time, the magnitude of these effects varied across the cohort, reflecting differences in individual immune reactivity. Further studies in larger cohorts will be valuable for characterizing this variability and identifying factors that may contribute to differences in vaccine induced immunity across populations. Furthermore, the immunological analyses presented here provide a preliminary characterization of these responses, and the limited number of flow cytometry time points does not permit a detailed assessment of their kinetics or persistence. Future longitudinal studies will therefore be important for determining the duration of protection and defining the development and maintenance of immunological memory.

Going forward, it is likely that the levels of Mtb protection can be improved through further optimization of the amounts, ratios, and dosing schedules of the MS2-H1_y_/MS2- cGAMP vaccine. Additionally, there is ongoing work toward engineering capsids that can carry both H1 and cGAMP on the same capsid, which may perform differently from the combination system reported herein. Lastly, to our knowledge all effective Mtb vaccines currently being investigated require three doses for robust protection. It is possible that the greater uptake levels of the MS2-H1/MS2-cGAMP platform will allow improved protection levels with fewer doses. Experiments to elucidate these points are currently underway. Taken together, these findings establish MS2-delivery of H1 and cGAMP as a promising platform for tuberculosis vaccination that has potential for human application. With further development, this approach could help bring the field one step closer to the control and eventual eradication of tuberculosis disease.

## METHODS

### Safety Comment

The synthesis of the linker for the attachment of cGAMP to MS2 involved two hazardous chemicals: phosgene and triethylammonium trihydrofluoride (Et_3_N•3 HF, HF-TEA). Phosgene is a toxic gas that should only be handled in a chemical fume hood and with PPE that includes nitrile gloves, a lab coat, and safety glasses. HF-TEA contains 37% HF, which causes severe burns and is toxic upon skin exposure. HF-TEA should only be used in a chemical fume hood with all PPE described for phosgene, in addition to using two pairs of nitrile gloves. Most importantly, always have calcium Calgonate gel close by in case of skin exposure to HF.

*Mycobacterium tuberculosis*, the causative agent of tuberculosis (TB), is an airborne pathogen that presents a significant risk of laboratory-acquired infection. Work involving viable *M. tuberculosis* should be conducted in a Biosafety Level 3 (BSL-3) laboratory in accordance with institutional biosafety policies, regulatory requirements, and approved risk assessments. Appropriate engineering controls, including specialized ventilation and containment equipment, should be used alongside suitable personal protective equipment (PPE) and personnel training. Procedures with the potential to generate aerosols require particular care and should be performed within certified biological safety cabinets or other approved containment devices. Established decontamination, waste management, and incident-reporting procedures must be followed to minimize the risk of exposure and maintain a safe working environment.

### Recombinant Expression and Purification of MS2 Mutants

MS2 mutants were expressed and purified as previously described.^13,17^ DH10B competent cells transformed with plasmids for each MS2 mutant were cultured overnight in 10 mL of Luria-Bertani (LB) broth with 25 µg/mL chloramphenicol for 16 h. The overnight cultures were inoculated into 1 L of 2x Yeast Extract Tryptone medium (2xYT) with 25 µg/mL chloramphenicol at 37 °C. To induce protein expression, arabinose was added to the flask at a final concentration of 0.1% w/v when optical density (OD_600_) measured ∼0.6 in approximately 3 h. After 22 h of growth at 37 °C, the cells were harvested.

To purify the MS2 mutants, cells were lysed via sonication in 10 mM sodium phosphate buffer with 2 mM sodium azide at pH 7.4 (SEC buffer) with 10 mM of DTT. After centrifugation, the supernatant was collected, and an equal volume of saturated ammonium sulfate was added to precipitate the capsids for 1 h. The precipitate was centrifuged at 14,000 rcf for 45 min, then resuspended in SEC buffer and filtered with a 0.2 µM sterile syringe filter. Next, four FPLC/Akta purification steps were performed. First, the filtrate was applied to two HiScreen Capto Core700 (Capto Core) columns in tandem using isocratic flow of SEC buffer, and the flowthrough was collected. The flowthrough from the Capto Core column was then desalted to remove excess ammonium sulfate using the HiPrep 26/10 Desalting column (Cytiva). Next, the HiPrep Heparin FF 16/10 column (Cytiva) was used to selectively bind MS2 mutants with KR mutations. The MS2 sample was applied to the heparin column using 20 mM sodium phosphate buffer, pH 7.5, then MS2 was eluted using a gradient from 0 to 100% of 20 mM sodium phosphate buffer with 2 M NaCl, pH 7.5. MS2s with KR mutations generally eluted in sodium phosphate buffer with approximately 1 M NaCl. Lastly, the MS2 fractions were desalted using the HiPrep Heparin FF 16/10 column (Cytiva). Final samples were concentrated and stored at 4 °C. MS2 samples were never frozen because of concerns with stability. MS2 variant masses were confirmed using ESI-TOF LC-MS and capsid assembly state was confirmed using HPLCSEC (described later). MS2 sequences are listed in SI Figure S1.

### Recombinant Expression, Insoluble Purification, TEV Cleavage, and Refolding of H1_y_ Antigen

H1-Tyr (H1_y_) was cloned into a pet-28b plasmid using Golden Gate assembly and transformed into One Shot™ BL21 Star™ (DE3) (Thermo Fisher) competent cells. A 1 L flask of LB was inoculated with 10 mL of overnight culture and kanamycin was added as a selection antibiotic. The 1 L culture was allowed to grow at 37 °C until OD_600_ = ∼0.6, then 1 mM IPTG was added to induce protein expression. After induction the culture was allowed to grow for 16 h at 25 °C then was harvested.

For the purification of H1_y_ from inclusion bodies, harvested cells were first resuspended in PBS + 0.1% Tween, pH 7.5 with 0.175 mg/mL PMSF protease inhibitor and lysed using sonication. The sonication pellet, which included inclusion bodies, was saved for the next steps. The pellet was washed three times with PBS + 0.1% Tween, pH 7.5, then washed an additional three times with PBS, pH 7.5. For each wash, the pellet was thoroughly vortexed in the buffer, then centrifuged at 14,000 rcf for 30 min. The washed pellet was then resuspended in denaturing equilibration buffer (PBS + 8 M urea + 10 mM imidazole, pH 7.5) on a rotator at 4 °C overnight. The resuspension was centrifuged at 14,000 rcf for 30 min and the supernatant was saved. Next, a 5 mL HisTrap HP (Cytiva) was equilibrated with denaturing equilibration buffer, and the resuspension supernatant was added to the column. H1_y_ was eluted using a gradient from the denaturing equilibration buffer to denaturing elution buffer (PBS + 8 M urea + 250 mM imidazole, pH 7.5). Next, H1_y_ was refolded to remove urea for TEV cleavage of the His_6_ tag. The His_6_ tag must be cleaved so H1_y_ does not bind to the nickel resin that megaTyr is bound to during the on-resin conjugation with MS2. To refold H1_y_, first, a Millipore Amicon Ultra Centrifugal Filter (10 kDa MWCO) was used to buffer exchange the protein into the solubilization buffer (50 mM CAPS buffer, 1.5% *N*-lauryl sarcosine, 300 mM NaCl, pH 11). H1_y_ was buffer exchanged into this buffer 5 times, with at least a 5× dilution each round. The resulting buffer exchanged H1 was then dialyzed into 2 L of PBS two times, for at least 8 h each time.

To TEV cleave the His_6_ tag from H1_y_, a 1:12 equivalent of TEV to H1 was added for 2 h with rotation at 4 °C. Because the H1 protein precipitated upon a standard nickel column purification, the TEV cleaved H1_y_ required a second denaturation for purification of H1_y_ from the His_6_ tag followed by refolding again. The denaturing 5 mL HisTrap HP column was performed the same as the previous nickel purification step, except the flowthrough containing cleaved H1_y_ was collected. The cleaved H1 was then refolded again as described in the previous paragraph. Proper expression of H1_y_ was confirmed using SDS-PAGE, with an expected mass of 43 kDa. H1_y_ was stored in PBS with 15 % v/v glycerol at −80 °C in aliquots until future use.

### Recombinant Expression of megaTyr Enzyme

The megaTyr tyrosinase enzyme was purified as described in a previous report.^29^ Briefly, a 10 mL starter culture was started using BL21(DE3) cells containing the megaTyr expression plasmid. This culture was used to inoculate 1 L of 2xYT media, and then 1 mM IPTG was used to induce protein expression at OD_600_ = ∼0.6. Cells were harvested after 16 h of growth at 37 °C, lysed using sonication, then purified using the 5 mL HisTrap HP (Cytiva) column. This HisTrap column was primed with 20 mM Tris-HCl + 500 mM NaCl + 10 mM imidazole, pH 7.5 and megaTyr was eluted with a gradient from 10 mM imidazole to 250 mM imidazole. The eluant was dialyzed into PBS with 20 µM copper(II) sulfate overnight at 4 °C. Protein mass was confirmed using ESI-TOF LC-MS and enzyme activity was measured as previously described.^29^ MegaTyr was stored at −80 °C in PBS with 20 µM copper(II) sulfate and 15% v/v glycerol.

### Binding of megaTyr Enzyme to Nickel Resin

A 50% slurry of HisPur Ni-NTA resin (Thermo Fisher) was centrifuged for 1 min at 13,000 rcf and the supernatant was removed. The resin was washed once using 100 mM sodium phosphate buffer, pH 8.0 then a second time using 100 mM sodium phosphate containing 300 mM NaCl and 10 mM imidazole, pH 8.0. Washed resin was suspended to approximately a 15% slurry in 100 mM sodium phosphate containing 300 mM NaCl and 10 mM imidazole, pH 8.0, and then 1,000 U/L megaTyr with a His_6_ tag was added to approximately a 1:1 v/v ratio of resin to megaTyr. The resin and megaTyr were co-incubated on a rotator at 4 °C for at least 1 h. Ni-megaTyr resin was then washed 3 times on a 0.2 µM spin filter: once with 100 mM sodium phosphate containing 300 mM NaCl and 10 mM imidazole, pH 8.0, and twice with 100 mM sodium phosphate buffer at pH 8.0. Each centrifugation was immediately followed by a resuspension of the resin in the next buffer to minimize the time the resin was dry. After three washes, the resin was suspended to a final ∼40-50% slurry (Ni-megaTyr resin) in 100 mM sodium phosphate buffer at pH 8.0. The pH of 8.0 was required to maintain strong binding of megaTyr to the resin. Ni-megaTyr resin must be made fresh and used day-of for acceptable enzymatic activity.

### Tyrosinase Enzymatic Coupling of H1_y_ to MS2

In 50 mM sodium phosphate buffer at pH 8.0, MS2 was added to a final concentration of 20 µM. The final concentration of H1_y_ in the reaction varied slightly, from 2 to 3 µM, depending on the activity of each specific batch of megaTyr on nickel resin. A ∼40-50% slurry of Ni-megaTyr resin was added to a final volume of 1/5 the final reaction volume, then the reaction was rotated at 4 °C for 90 min. MS2- H1_y_ conjugation reactions were completed at pH 8 to ensure the megaTyr enzyme remained strongly bound to the resin for complete removal by filtration when the reaction was complete.^13^ After 90 min, the reaction was filtered to remove Ni-megaTyr resin and buffer exchanged 7 times into DPBS at pH 7.5 (with 5× dilution each buffer exchange round). Reactions were analyzed for % modification of H1_y_ on MS2 using SDS-PAGE followed by densitometry using ImageJ.

Additionally, HPLC-SEC was used to analyze the capsid assembly state. Only reactions with one peak at the expected assembled MS2 elution time were used for vaccination.

The acceptable range of H1_y_ conjugation of MS2 was very narrow (8-13%). This narrow range was to ensure there was enough antigen for vaccine efficacy but to not destabilize the MS2 capsid. Because each batch of Ni-megaTyr resin must be made fresh and could vary slightly in activity, several small-scale reactions were performed the same day before each large-scale conjugation for use in vaccination. The small-scale reactions were used to estimate the most effective stoichiometric ratio of H1_y_ to use for each batch, which ranged from 2 to 3 µM.

### Recombinant Expression of cGAS Enzyme

cGAS was cloned with a SUMO tag for soluble protein expression. The enzyme was expressed from a 10 mL overnight culture in LB, inoculated into 1 L of terrific broth. The culture was allowed to grow at 37 °C until OD_600_ = ∼0.6, then was induced with 0.5 mM IPTG. The temperature was lowered to 18 °C for protein expression, and cells were harvested 16 h later.

cGAS purification must be done in one day. To purify cGAS, the cells were lysed using sonication in cGAS equilibration buffer (50 mM Tris + 300 mM NaCl + 20 mM Imidazole + 1 mM freshly added TCEP, pH 8.0) with 0.175 mg/mL PMSF protease inhibitor. All lines of the Akta FPLC must be washed thoroughly prior to cGAS purification to ensure cGAS does not precipitate due to contaminants on the lines. Each line was washed thoroughly at 5mL/min in this order: Milli-Q water, 70% Acetic acid, Milli-Q water, 2 M NaOH, Milli-Q water. A 5 mL HisTrap HP (Cytiva) column was used to purify the SUMO-cGAS fusion using a gradient from the cGAS equilibration buffer to the cGAS elution buffer (50 mM Tris + 300 mM NaCl + 300 mM imidazole + 1 mM fresh TCEP, pH 8.0). Eluted cGAS was immediately desalted into cGAS desalting buffer (50 mM Tris + 1 mM fresh TCEP) using a HiPrep 26/10 Desalting column (Cytiva).

### Enzymatic Synthesis of cGAMP

cGAS was used for enzymatic cGAMP synthesis immediately following purification. Together in one flask, 50 mM Tris, pH 8.0 was used to dissolve each component to their given concentration: 1 mM TCEP-HCl, 10 mM magnesium chloride, 2 mM ATP, 2 mM GTP, 0.05 mg/mL DNA (Sigma-Aldrich D9156), 1 µM cGAS. This reaction was stirred at 37

°C for 24 h. The complete reaction was opaque white from cGAS precipitation over time. Upon reaction completion, the majority of the buffer was removed using a rotary evaporator until between 10 and 30 mL remained. Remaining solution was frozen at −80 °C then lyophilized.

Lyophilized product was resuspended in 10 mL of milli-Q water and centrifuged at 14,000 rcf for 10 min to remove precipitated cGAS. The supernatant was wet loaded onto a Teledyne ISCO RediSep^®^Rf Gold 15.5 g C18 column using an Isolera One ACI™ Biotage and was purified using a reverse phase gradient from 50 mM TEAA buffer in H_2_O to acetonitrile. The acetonitrile was removed using rotary evaporation, then the product was lyophilized. Synthesis of cGAMP was confirmed using small molecule ESI-TOF LCMS.

### Chemical Synthesis Self-Immolative Disulfide linker

The self-immolative linker was synthesized as previously described.^16^ Briefly, in a round bottom flask, 2 g of 2,2′-dipyridyl disulfide was dissolved in 25 mL of methanol with 1.5 mL glacial acetic acid. Separately, 10 mL of methanol was added to a separatory funnel and 2-mercapto ethanol (BME) at a 2:1 molar ratio of 2,2′-dipyridyl disulfide to BME. The BME in methanol was mixed in the addition funnel, then added dropwise to the reaction flask. The reaction was stirred overnight at RT. The reaction turned bright yellow, signaling successful disulfide exchange. TLC using 50/50 hexanes to ethyl acetate (EtOAc) was used to confirm reaction completion, where the desired product was the middle of three spots. The crude product was a light-yellow oil. The linker product was purified twice using two silica flash columns (Teledyne ISCO Flash Column (80g), followed by Teledyne ISCO RediSep^®^ Silver 12g Silica) using a normal phase solvent system of hexanes and EtOAc. The final pure product was a very pale-yellow oil that formed a white solid upon freezing at −20 °C.

### Addition of Linker to cGAMP

cGAMP was modified with the disulfide linker using a modified version of a previously published protocol.^16^ To activate the linker for cGAMP modification, the disulfide linker was added to a round bottom flask, and the flask was sealed then purged using three rounds of alternating vacuum and nitrogen gas on a Schlenk line. Dry dichloromethane (DCM) was added, and the flask was cooled to −78 °C using dry ice and acetone. Next, dry phosgene was added dropwise at a 30:100 ratio of linker to phosgene and the reaction was stirred at −78 °C for 1 h under nitrogen. Upon reaction completion, volatiles were removed *in vacuo*, and the product was azeotroped three times with dry DCM to remove excess phosgene. In a separate vial, cGAMP was weighed and suspended using dry pyridine (a final ratio of 30:1 linker to cGAMP was used). Triethylammonium trihydrofluoride (Et_3_N•3 HF, HF-TEA, Thermo Scientific Cat. No: L14417.06) was added to the vial with cGAMP and pyridine until all visible cGAMP dissolved using a water bath sonicator (approximately 200 µL of HF-TEA per 10 mg of cGAMP). The dissolved cGAMP with HF-TEA was added to the reaction flask and stirred at room temperature (RT) for 90 min. Volatiles were removed *in vacuo* and the product was frozen at −20 °C until purification. The cGAMP-linker product was purified using the same reverse phase method as described for cGAMP purification, using an ISCO RediSep^®^Rf Gold 15.5 g C18 column. The final product was lyophilized stored at −20 °C.

### Conjugation of cGAMP to MS2

To conjugate MS2 to cGAMP, MS2 PKR was diluted to a 50 µM concentration in SEC buffer. cGAMP-linker was added at a 1:5 ratio of MS2 to cGAMP-linker and allowed to react at 4 °C overnight. Excess cGAMP-linker was removed using 7 rounds of buffer exchange in a 100 kDa MWCO centrifugal filter (Millipore), with 5-fold dilution each round. The product MS2-cGAMP was verified using ESI-TOF LC-MS and capsid assembly was confirmed using HPLC-SEC and TEM as described below.

### Ethics Statement for *In Vivo* Protocols

All procedures involving the use of mice were approved by the University of California, Berkeley Institutional Animal Care and Use Committee (AUP20200713458). All protocols conform to federal regulations, the National Research Council Guide for the Care and Use of Laboratory Animals, and the Public Health Service Policy on Humane Care and Use of Laboratory Animals.

### Vaccine Reagents

ML-RR-cGAMP was synthesized at Aduro Biotech as described previously.^37^ H1 fusion protein was provided by Aeras and Statens Serum Institute, and peptide pools were provided by the NIH BEI Resources Repository. Vaccine combinations were made in PBS. Production of MS2, H1_y_, and cGAMP, and conjugation of MS2 to cGAMP and H1_y_, was conjugated to cGAMP and H1 as described above.

### Mice

C57BL/6J (#000664) mice were obtained from The Jackson Laboratory (Bar Harbor, ME). Mice were sex- and age-matched for all experiments.

### Mtb Bacterial Culture

*M. tuberculosis* strain Erdman was grown in Middlebrook 7H9 liquid medium supplemented with 10% oleic acid, albumin, dextrose, catalase (OADC), 0.4% glycerol, and 0.05% Tween 80 or on solid 7H10 agar plates supplemented with 10% Middlebrook OADC (BD Biosciences) and 0.4% glycerol. Frozen stocks of M. tuberculosis were made from single cultures and used for all experiments.

### Vaccinations

Mice were vaccinated three times at 4-week intervals with 20 μL *i*.*n*. of either PBS, 5 μg ML-RR-cGAMP (CDN) and 3 μg fusion antigen protein (H1), 86 ng MS2-cGAMP and 1 μg MS2-H1_y_, 86 ng cGAMP and 1μg H1, 1 μg MS2-H1_y_, 86 ng MS2-cGAMP, 33.6 ng MS2-cGAMP and 1 μg MS2-H1_y_, 33.6 ng cGAMP and 1 μg H1, 33.6 ng MS2-cGAMP, 86 ng MS2- cGAMP and 0.53 μg MS2-H1_y_. All mass values are given with respect to the mass of cGAMP and H1, not the mass of the total MS2 conjugate. All vaccines were formulated in PBS for intranasal (*i*.*n*.) delivery.

### Challenge Experiments with *M. tuberculosis*

Twelve weeks after the initial vaccine injection, mice were infected via the aerosol route with *M. tuberculosis* Erdman strain. Aerosol infection was done using a nebulizer and fullbody inhalation exposure system (Glas-Col, Terre Haute, IN). A total of 9 mL of culture diluted in sterile water was loaded into the nebulizer calibrated to deliver ∼30 bacteria per mouse as measured by CFU in the lungs 1 day following infection (data not shown).

### Pre-challenge ICS Assay

Heparinized blood lymphocytes were isolated 9 weeks post priming. Blood was collected by retro-orbital bleeds into heparinized tubes and processed using Red Blood Lysis buffer followed by a series of washes in cRPMI. Cell suspensions were pelleted and resuspended in 400 μL of cRPMI. A 100 uL sample of cells was restimulated with either no peptide, Ag85B, or ESAT6 peptide pools (2 μg/mL) with Protein Transport Inhibitor Cocktail (eBioscience 00-4980-93) for 5 h at 37 °C. Cells were washed with FACS buffer and stained with Live/Dead stain (Thermo Fisher Scientific L34970), Fc receptor block, and antibodies specific for CXCR3, major histocompatibility complex (MHC) class II, TCRγ/δ, CD8a, IFNg, and IL-17 (BioLegend 101320, 126522, 107606, 118124, and 100750 respectively), and CD4, CD90.2, KLRG1 and Ly6G (BD Biosciences 564933, 561616, 740279 and 551460, respectively), and fixed/permeabilized with BD Cytofix/Cytoperm Fixation/Permeabilization Solution Kit (Thermo Fisher, 554714) before staining with antibodies specific for IFN-γ (BioLegend 505816) and IL-17 (BioLegend 506904). Data were collected using a BD LSR Fortessa flow cytometer and analyzed using FlowJo Software (Tree Star, Ashland, OR).

### Post-Challenge Tissue Processing for Bacterial Enumeration and Flow Cytometry

Mice were harvested 28 days post-infection to measure CFUs by plating and immune cell populations by flow cytometry. All lung lobes were harvested into a gentleMACS C tube (Miltenyi Biotec) containing 3 mL of RPMI media with 70 μg / mL of Liberase TM (Roche) and 30 μg / mL of Dnase I (Roche). Samples were processed into chunks using the lung_01 setting on the gentleMACS (Miltenyi Biotec) and incubated for 30 min at 37 °C. Tissue was then homogenized into a single cell suspension by running the samples on the lung_02 setting on the gentleMACS. The digestion was quenched by adding 2 mL of PBS with 20% Newborn Calf Serum (Thermo Fisher Scientific) and filtered through 70 μm SmartStrainers (Miltenyi Biotec).

For bacterial enumeration, 100 μL was taken from each single cell suspension and then serially diluted in phosphate-buffered saline (PBS) with 0.05% Tween-80. Serial dilutions were plated on 7H10 plates and colonies were counted 3 weeks after plating.

For flow cytometry, single lung cell suspensions were pelleted and resuspended in 400 μL of cRPMI. Samples (100 µL) of cells were restimulated, washed, and stained with antibodies as described in the pre-challenge ICS. Data were collected and analyzed as outlined above.

### Statistical Analysis

Data are presented as mean values, and error bars represent SD. Symbols represent individual animals. The number of samples and statistical tests used are denoted in the legend of the corresponding figure for each experiment. Analysis of statistical significance was performed using GraphPad Prism 8 (GraphPad, La Jolla, CA), and p < 0.05 was considered significant.

## ASSOCIATED CONTENT

### Supporting Information

Materials, analytical methods, equipment, protein and peptide sequences and properties, and supplementary figures. The Supporting Information is available free of charge on the ACS Publications website.

## AUTHOR CONTRIBUTIONS

The manuscript was written through contributions of all authors. All authors have given approval to the final version of the manuscript.

### Notes

No notes to add.

## ACKNOWLEDGMENT

We would like to thank the David Raulet lab at UC Berkeley for sharing their THP-1 STING reporter cell line. We would like to thank the staff at the University of California Berkeley Electron Microscope laboratory for their advice and assistance in electron microscopy sample preparation and data collection.

## FUNDING SOURCES

Funding for this work was provided by the UCSF Center for Tuberculosis, NIH/NIAID P30: TB Research Advancement Center (UC TRAC), P30AI168440 and NIH/NIAID R25: TB Research and Mentorship Program (TB RAMP), 1R25AI147375. Additionally, this material is based upon work supported by the National Science Foundation Graduate Research Fellowship Program under Grant No. DGE 2146752 for HSM. Any opinions, findings, and conclusions or recommendations expressed in this material are those of the author(s) and do not necessarily reflect the views of the National Science Foundation.

## CONFLICT OF INTEREST

MBF has a financial interest in Catena Bio-sciences, which has licensed the technology used to make the conjugates described herein, and both he and the company may benefit from commercialization of the results of this research. SAS and MBF are strategic advisors for X-biotix Therapeutics. All other authors declare no competing interests.

### For Table of Contents only

**Synopsis:**
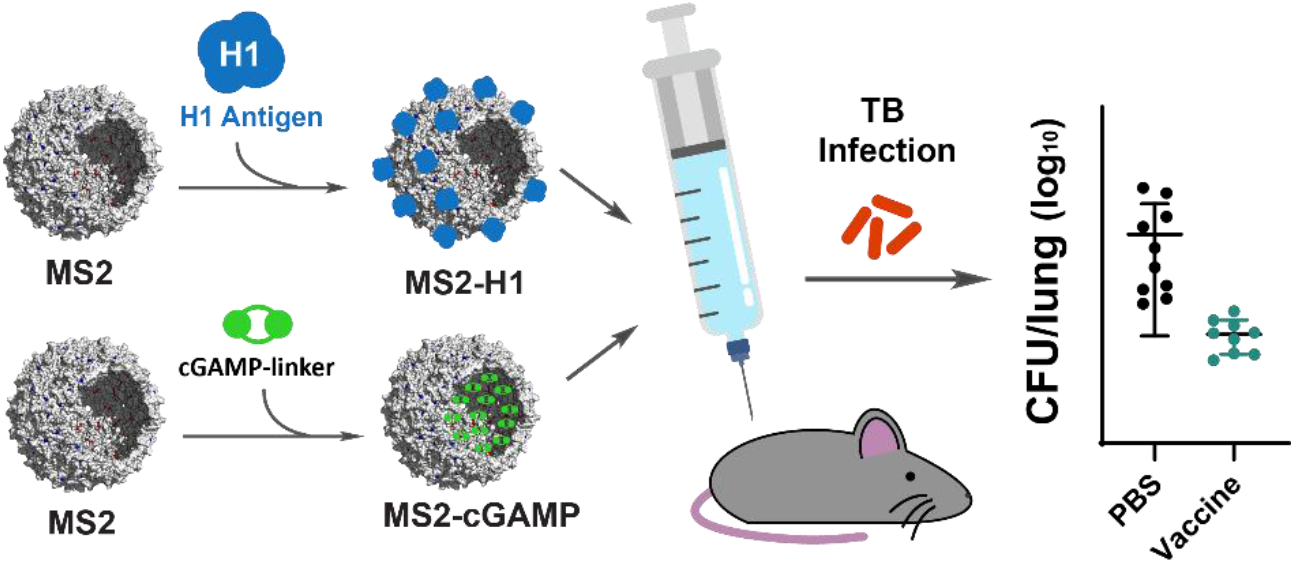
An MS2 viral capsid-delivered vaccine provided protection against tuberculosis infection with 57-fold less adjuvant and 3-fold less antigen compared to a previously protective vaccine, increasing potency and lowering cost.

## Notes

### Summary of Updates

The manuscript has been revised to include a couple brief explanations of the limitations to the immunological experiments conducted in this work. Additionally, one additional figure was added to the supporting information. This figure includes further controls (two doses of MS2-cGAMP only) for the in vivo experiment that showed protection against Mtb infection. These controls prove that each component of the vaccine, MS2-H1 and MS2-cGAMP, are required for protection. Additionally, this new figure demonstrated equal to greater protection against infection at an even lower dose of MS2-cGAMP in the vaccine; this dose used 145-fold less STING agonist adjuvant than the previously established H1/CDN vaccine.

## REFERENCES

(1) Loddenkemper, R.; Murray, J. F. History of Tuberculosis. Essential Tuberculosis 2021, 3–9. 10.1007/978-3-030-66703-0_1.

(2) Barberis, I.; Bragazzi, N. L.; Galluzzo, L.; Martini, M. The History of Tuberculosis: From the First Historical Records to the Isolation of Koch’s Bacillus. J. Prev. Med. Hyg. 2017, 58 (1), E9.

(3) Andersen, P.; Doherty, T. M. The Success and Failure of BCG — Implications for a Novel Tuberculosis Vaccine. Nature Reviews Microbiology 2005 3:8 2005, 3 (8), 656–662. 10.1038/nrmicro1211.

(4) Mangtani, P.; Abubakar, I.; Ariti, C.; Beynon, R.; Pimpin, L.; Fine, P. E. M.; Rodrigues, L. C.; Smith, P. G.; Lipman, M.; Whiting, P. F.; Sterne, J. A. Protection by BCG Vaccine Against Tuberculosis: A Systematic Review of Randomized Controlled Trials. Clinical Infectious Diseases 2014, 58 (4), 470–480. 10.1093/CID/CIT790.

(5) Jong, R. M.; Van Dis, E.; Berry, S. B.; Nguyenla, X.; Baltodano, A.; Pastenkos, G.; Xu, C.; Fox, D.; Yosef, N.; McWhirter, S. M.; Stanley, S. A. Mucosal Vaccination with Cyclic Dinucleotide Adjuvants Induces Effective T Cell Homing and IL-17–Dependent Protection against Mycobacterium Tuberculosis Infection. The Journal of Immunology 2022, 208 (2), 407–419. 10.4049/JIMMUNOL.2100029.

(6) Van Dis, E.; Sogi, K. M.; Rae, C. S.; Sivick, K. E.; Surh, N. H.; Leong, M. L.; Kanne, D. B.; Metchette, K.; Leong, J. J.; Bruml, J. R.; Chen, V.; Heydari, K.; Cadieux, N.; Evans, T.; McWhirter, S. M.; Dubensky, T. W.; Portnoy, D. A.; Stanley, S. A. STING-Activating Adjuvants Elicit a Th17 Immune Response and Protect against Mycobacterium Tuberculosis Infection. Cell Rep. 2018, 23 (5), 1435. 10.1016/J.CELREP.2018.04.003.

(7) Decout, A.; Katz, J. D.; Venkatraman, S.; Ablasser, A. The CGAS–STING Pathway as a Therapeutic Target in Inflammatory Diseases. Nature Reviews Immunology 2021 21:9 2021, 21 (9), 548–569. 10.1038/s41577-021-00524-z.

(8) Pan, J.; Fei, C. J.; Hu, Y.; Wu, X. Y.; Nie, L.; Chen, J. Current Understanding of the CGAS-STING Signaling Pathway: Structure, Regulatory Mechanisms, and Related Diseases. Zool. Res. 2023, 44 (1), 183. 10.24272/J.ISSN.2095-8137.2022.464.

(9) Meric-Bernstam, F.; Sweis, R. F.; Hodi, F. S.; Messersmith, W. A.; Andtbacka, R. H. I.; Ingham, M.; Lewis, N.; Chen, X.; Pelletier, M.; Chen, X.; Wu, J.; McWhirter, S. M.; Müller, T.; Nair, N.; Luke, J. J. Phase I Dose-Escalation Trial of MIW815 (ADU-S100), an Intratumoral STING Agonist, in Patients with Advanced/Metastatic Solid Tumors or Lymphomas. Clinical Cancer Research 2022, 28 (4), 677–688. 10.1158/1078-0432.CCR-21-1963/674040/AM/PHASE-I-DOSE-ESCALATION-TRIAL-OF-MIW815-ADU-S100.

(10) Dosta, P.; Cryer, A. M.; Prado, M.; Artzi, N. Bioengineering Strategies to Optimize STING Agonist Therapy. Nature Reviews Bioengineering 2025 3:8 2025, 3 (8), 660–680. 10.1038/s44222-025-00337-y.

(11) Valegard, K.; Liljas, L.; Fridborg, K.; Unge, T. The Three-Dimensional Structure of the Bacterial Virus MS2. Nature 1990, 345, 36–41.

(12) Golmohammadi, R.; Valegård, K.; Fridborg, K.; Liljas, L. The Refined Structure of Bacteriophage MS2 at 2·8 Å Resolution. J. Mol. Biol. 1993, 234 (3), 620–639. 10.1006/JMBI.1993.1616.

(13) Martin, H. S.; Huang, P.; Leifer, I. C.; Pratakshya, P.; Francis, M. B. Engineered MS2 Virus Capsids for Cellular Display of Peptide Antigens. ACS Chem. Biol. 2025, 20 (12), 2943–2954. 10.1021/ACSCHEMBIO.5C00700.

(14) Huang, P.; Jo, Y.; Martin, H. S.; Luteijn, R. D.; Raulet, D. H.; Francis, M. B. Viral Capsid Delivery of CGAMP Enhances STING-Dependent Antitumor Immune Response. bioRxiv 2026, 2026.06.26.734859. 10.64898/2026.06.26.734859.

(15) Pistono, P. E.; Huang, P.; Brauer, D. D.; Francis, M. B. Fitness Landscape-Guided Engineering of Locally Supercharged Virus-like Particles with Enhanced Cell Uptake Properties. ACS Chem. Biol. 2022, 17 (12), 3367–3378. 10.1021/ACSCHEMBIO.2C00318.

(16) Huang, P.; Pistono, P. E.; Martin, H. S.; Fetzer, J. L.; Francis, M. B. Cytosolic Delivery of Anionic Cyclic Dinucleotide STING Agonists with Locally Supercharged Viral Capsids. ACS Chem. Biol. 2025, 20 (8), 1875–1883. 10.1021/ACSCHEMBIO.5C00125.

(17) Pistono, P. E.; Xu, J.; Huang, P.; Fetzer, J. L.; Francis, M. B. Exploring the Effects of Intersubunit Interface Mutations on Virus-Like Particle Structure and Stability. Biochemistry 2024, 63 (15), 1913–1924. 10.1021/ACS.BIOCHEM.4C00225.

(18) Tsolaki, A. G.; Nagy, J.; Leiva, S.; Kishore, U.; Rosenkrands, I.; Robertson, B. D. Mycobacterium Tuberculosis Antigen 85B and ESAT-6 Expressed as a Recombinant Fusion Protein in Mycobacterium Smegmatis Elicits Cell-Mediated Immune Response in a Murine Vaccination Model. Mol. Immunol. 2013, 54 (3–4), 278–283. 10.1016/J.MOLIMM.2012.11.014.

(19) Khoshnood, S.; Heidary, M.; Haeili, M.; Drancourt, M.; Darban-Sarokhalil, D.; Nasiri, M. J.; Lohrasbi, V. Novel Vaccine Candidates against Mycobacterium Tuberculosis. Int. J. Biol. Macromol. 2018, 120, 180–188. 10.1016/J.IJBIOMAC.2018.08.037.

(20) Anderson, D. H.; Harth, G.; Horwitz, M. A.; Eisenberg, D. An Interfacial Mechanism and a Class of Inhibitors Inferred from Two Crystal Structures of the Mycobacterium Tuberculosis 30 Kda Major Secretory Protein (Antigen 85B), a Mycolyl Transferase. J. Mol. Biol. 2001, 307 (2), 671–681. 10.1006/JMBI.2001.4461.

(21) Moguche, A. O.; Musvosvi, M.; Penn-Nicholson, A.; Plumlee, C. R.; Mearns, H.; Geldenhuys, H.; Smit, E.; Abrahams, D.; Rozot, V.; Dintwe, O.; Hoff, S. T.; Kromann, I.; Ruhwald, M.; Bang, P.; Larson, R. P.; Shafiani, S.; Ma, S.; Sherman, D. R.; Sette, A.; Lindestam Arlehamn, C. S.; McKinney, D. M.; Maecker, H.; Hanekom, W. A.; Hatherill, M.; Andersen, P.; Scriba, T. J.; Urdahl, K. B. Antigen Availability Shapes T Cell Differentiation and Function during Tuberculosis. Cell Host Microbe 2017, 21 (6), 695. 10.1016/J.CHOM.2017.05.012.

(22) Sefat, K. M. S. R.; Kumar, M.; Kehl, S.; Kulkarni, R.; Leekha, A.; Paniagua, M. M.; Ackart, D. F.; Jones, N.; Spencer, C.; Podell, B. K.; Ouellet, H.; Varadarajan, N. An Intranasal Nanoparticle Vaccine Elicits Protective Immunity against Mycobacterium Tuberculosis. Vaccine 2024, 42 (22), 125909. 10.1016/J.VACCINE.2024.04.055.

(23) Malik, A.; Gupta, M.; Mani, R.; Bhatnagar, R. Single-Dose Ag85B-ESAT6–Loaded Poly(Lactic-Co-Glycolic Acid) Nanoparticles Confer Protective Immunity against Tuberculosis. Int. J. Nanomedicine 2019, 14, 3129. 10.2147/IJN.S172391.

(24) Foreman, T. W.; Mehra, S.; Lackner, A. A.; Kaushal, D. Translational Research in the Nonhuman Primate Model of Tuberculosis. ILAR J. 2017, 58 (2), 151–159. 10.1093/ILAR/ILX015.

(25) Agger, E. M.; Rosenkrands, I.; Olsen, A. W.; Hatch, G.; Williams, A.; Kritsch, C.; Lingnau, K.; von Gabain, A.; Andersen, C. S.; Korsholm, K. S.; Andersen, P. Protective Immunity to Tuberculosis with Ag85B-ESAT-6 in a Synthetic Cationic Adjuvant System IC31. Vaccine 2006, 24 (26), 5452–5460. 10.1016/J.VACCINE.2006.03.072.

(26) Olsen, A. W.; Van Pinxteren, L. A. H.; Okkels, L. M.; Rasmussen, P. B.; Andersen, P. Protection of Mice with a Tuberculosis Subunit Vaccine Based on a Fusion Protein of Antigen 85B and ESAT-6. Infect. Immun. 2001, 69 (5), 2773–2778. 10.1128/IAI.69.5.2773-2778.2001;WGROUP:STRING:PUBLICATION.

(27) Dietrich, J.; Andersen, C.; Rappuoli, R.; Doherty, T. M.; Jensen, C. G.; Andersen, P. Mucosal Administration of Ag85B-ESAT-6 Protects against Infection with Mycobacterium Tuberculosis and Boosts Prior Bacillus Calmette-Guérin Immunity. The Journal of Immunology 2006, 177 (9), 6353–6360. 10.4049/JIMMUNOL.177.9.6353.

(28) Lobba, M. J.; Fellmann, C.; Marmelstein, A. M.; Maza, J. C.; Kissman, E. N.; Robinson, S. A.; Staahl, B. T.; Urnes, C.; Lew, R. J.; Mogilevsky, C. S.; Doudna, J. A.; Francis, M. B. Site-Specific Bioconjugation through Enzyme-Catalyzed Tyrosine–Cysteine Bond Formation. ACS Cent. Sci. 2020, 6 (9), 1564–1571. 10.1021/ACSCENTSCI.0C00940.

(29) Marmelstein, A. M.; Lobba, M. J.; Mogilevsky, C. S.; Maza, J. C.; Brauer, D. D.; Francis, M. B. Tyrosinase-Mediated Oxidative Coupling of Tyrosine Tags on Peptides and Proteins. J. Am. Chem. Soc. 2020, 142 (11), 5078–5086. 10.1021/JACS.9B12002.

(30) Valenti, L. E.; De Pauli, C. P.; Giacomelli, C. E. The Binding of Ni(II) Ions to Hexahistidine as a Model System of the Interaction between Nickel and His-Tagged Proteins. J. Inorg. Biochem. 2006, 100 (2), 192–200. 10.1016/J.JINORGBIO.2005.11.003.

(31) Blest, H. T. W.; Chauveau, L. CGAMP the Travelling Messenger. Front. Immunol. 2023, 14, 1150705. 10.3389/FIMMU.2023.1150705.

(32) Luteijn, R. D.; Zaver, S. A.; Gowen, B. G.; Wyman, S. K.; Garelis, N. E.; Onia, L.; McWhirter, S. M.; Katibah, G. E.; Corn, J. E.; Woodward, J. J.; Raulet, D. H. SLC19A1 Transports Immunoreactive Cyclic Dinucleotides. Nature 2019, 573 (7774), 434–438. 10.1038/S41586-019-1553-0;TECHMETA.

(33) Bharadwaj, R.; Jaiswal, S.; Silverman, N. Cytosolic Delivery of Innate Immune Agonists. Trends Immunol. 2024, 45 (12), 1001–1014. 10.1016/j.it.2024.10.007.

(34) Kulkarni, R.; Maranholkar, V.; Nguyen, N.; Cirino, P. C.; Willson, R. C.; Varadarajan, N. The Efficient Synthesis and Purification of 2′3’-CGAMP from Escherichia Coli. Front. Microbiol. 2024, 15, 1345617. 10.3389/FMICB.2024.1345617/FULL.

(35) Kranzusch, P. J.; Lee, A. S. Y.; Berger, J. M.; Doudna, J. A. Structure of Human CGAS Reveals a Conserved Family of Second-Messenger Enzymes in Innate Immunity. Cell Rep. 2013, 3 (5), 1362–1368. 10.1016/j.celrep.2013.05.008.

(36) Sakai, S.; Kauffman, K. D.; Schenkel, J. M.; McBerry, C. C.; Mayer-Barber, K. D.; Masopust, D.; Barber, D. L. Cutting Edge: Control of Mycobacterium Tuberculosis Infection by a Subset of Lung Parenchyma-Homing CD4 T Cells. J. Immunol. 2014, 192 (7), 2965–2969. 10.4049/JIMMUNOL.1400019.

(37) Gaffney, B. L.; Veliath, E.; Zhao, J.; Jones, R. A. One-Flask Syntheses of c-Di-GMP and the [Rp,Rp] and [Rp,Sp] Thiophosphate Analogues. Org. Lett. 2010, 12 (14), 3269–3271. 10.1021/OL101236B.

