## Supporting Information for "Enhancement of a STING Agonist Vaccine for Tuberculosis Using Locally Supercharged MS2 Viral Capsids"

**Table of Contents**

**Materials…………………………………………………………………………………………………………..…2**

**Analytical Instrument Methods………………………………………………………………………………….…2**

**Supplementary Figures……………………………………………………………………………………………3-8**

**Materials**

**General Reagents**

Cloning: DNA primers were purchased from Integrated DNA Technologies. All golden gate assembly enzymes and components were purchased from New England BioLabs, and the Wizard® SV Gel and PCR CleanUp System was purchased from Promega. The ZymoPURE^TM^ Plasmid miniprep kit was purchased from Zymo Research. DNA sequencing was performed by Genewiz from Azenta and the UC Berkeley DNA sequencing facility.

Protein buffers and enzymatic cGAMP synthesis: Monobasic sodium phosphate and sodium chloride were purchased from thermo fisher, dibasic sodium phosphate was purchased from Spectrum Chemical MFG CORP. TWEEN 20 was purchased from Sigma-Aldrich (SigmaUltra), Sodium Lauroyl Sarcosine was from bioWORLD. ATP and GTP were purchased from Thermo Scientific.

**Cell Lines and Cell Culture Reagents**

THP-1 STING reporter cells were engineered by the Raulet lab at UC Berkeley and were generously shared with us. The THP-1 STING reporter cells were cultured in RPMI + 10% FBS at 37 °C with 5% CO_2_. RPMI and DPBS were purchased from Gibco. FBS was purchased from Avantor (Avantor Seradigm Premium Grade FBS, 100% United States Origin).

All other materials, equipment, and regents are specified within the methods section of the main text and instrument methods in the SI.

**Analytical Instrument Methods**

**Mass Spectrometry (LC-MS TOF) Analysis**

Mass spectrometry was used to confirm molecular weights of expressed proteins, small molecules, and to confirm modification of cGAMP on MS2. The LC-MS system used was an Agilent 1260 series liquid chromatograph and an Agilent 650 LC/QTOF mass spectrometer with electrospray ionization. For each protein analysis, the sample was injected into a Proswift RP-4H monolithic analytical column using a mobile phase of milli-Q water with 0.1% (v/v) MS-grade formic acid and MS grade MeCN (Fisher Optima) with 0.1% (v/v) MS-grade formic acid. For small molecule mass spectrometry, a WatersTM Acquity UPLC BEH C18 column was used with the same solvent and LC/QTOF system. Protein LC-MS deconvolution was completed using Mass Hunter BioConfirm Software 10.0. Data were graphed using Chartograph software (<http://chartograph.com>).

**HPLC-SEC Analysis of Conjugates**

HPLC-SEC was used to analyze all MS2 capsid expressions and MS2 vaccine conjugates to measure the capsid assembly state. Samples were injected into an Agilent Technologies 1260 infinity HPLC with an Agilent Bio SEC-5 column. The SEC method involved isocratic flow of 10 mM sodium phosphate buffer with 2 mM sodium azide at pH 7.4. Tryptophan fluorescence was measured with 280 nm excitation and 350 nm emission. An elution time of ~7 min corresponded to assembled MS2 capsids, whereas elution times of ~12 min or ~16-18 min corresponded to dissembled oligomers and monomers, respectively.

**TEM Analysis of Conjugates**

Formvar/carbon-coated copper 400 mesh grids (Electron Microscopy Sciences) were glow-discharged immediately prior to use. Subsequently, MS2 samples (5 µL, ~50 µM) were pipetted onto the grids for 2 min, followed by three washes with MilliQ water. Excess liquid was blotted with Whatman filter paper, after which 5 µL of 1% (w/v) uranyl acetate was applied for 2 min as a negative stain. Residual stain was blotted using a Whatman filter paper, and grids were air-dried under ambient conditions. Samples were imaged on a Tecnai 12 transmission electron microscope equipped with a 2k × 2k CCD camera at magnifications of 23,000×, 30,000×, and 49,000×. All TEM measurements were carried out at the Electron Microscope Lab at UC Berkeley.

**DLS Measurements of Conjugates**

A Malvern Zetasizer Nano-ZS DLS was used to measure capsid sizes. To a cuvette, 100 µL of a 50 µM protein sample was added with careful attention to avoid bubble formation. Three replicate number distribution measurements were recorded to calculate the mean number and standard deviation of capsid diameters.

**Supplementary Figures:**

**Proteins**

| **Protein** | **Abbreviation** | **Sequence** | **Molecular Weight** |
| --- | --- | --- | --- |
| MS2 **T71K G73R S120C** N87C | **MS2 KR S120C** | ASNFTQFVLVDNGGTGDVTVAPSNFANGVAEWISSNSRSQAYKVTCSVRQSSAQNRKYTIKVEVPKVATQKVRGVELPVAAWRSYLNMELTIPIFATNSDCELIVKAMQGLLKDGNPIPCAIAANSGIY | 13,870 |
| MS2 **S37P T71K G73R** N87C | **MS2 PKR** | ASNFTQFVLVDNGGTGDVTVAPSNFANGVAEWISSN**P**RSQAYKVTCSVRQSSAQNRKYTIKVEVPKVATQKVRGVELPVAAWRSYLCMELTIPIFATNSDCELIVKAMQGLLKDGNPIPSAIAANSGIY | 13,854 |
| H1-Tyr  (His12 tag-TEV cleavage site-H1-SG4-**Y**) | **H1_y_** | MHHHHHHHHHHHHENLYFQGSIEGRSFSRPGLPVEYLQVPSPSMGRDIKVQFQSGGNNSPAVYLLDGLRAQDDYNGWDINTPAFEWYYQSGLSIVMPVGGQSSFYSDWYSPACGKAGCQTYKWETFLTSELPQWLSANRAVKPTGSAAIGLSMAGSSAMILAAYHPQQFIYAGSLSALLDPSQGMGPSLIGLAMGDAGGYKAADMWGPSSDPAWERNDPTQQIPKLVANNTRLWVYCGNGTPNELGGANIPAEFLENFVRSSNLKFQDAYNAAGGHNAVFNFPPNGTHSWEYWGAQLNAMKGDLQSSLGAGKLAMTEQQWNFAGIEAAASAIQGNVTSIHSLLDEGKQSLTKLAAAWGGSGSEAYQGVQQKWDATATELNNALQNLARTISEAGQAMASTEGNVTGMFAGGSGGGGSGGGGSGGGGSGGGG**Y** | 45,657 (not TEV cleaved)  Or  43,029 (TEV cleaved) |
| Wild-type megaTyr  (tyrosinase from *Bacillus megaterium*) | **megaTyr** | SNKYRVRKNVLHLTDTEKRDFVRTVLILKEKGIYDRYIAWHGAAGKFHTPPGSDRNAAHMSSAFLPWHREYLLRFERDLQSINPEVTLPYWEWETDAQMQDPSQSQIWSADFMGGNGNPIKDFIVDTGPFAAGRWTTIDEQGNPSGGLKRNFGATKEAPTLPTRDDVLNALKITQYDTPPWDMTSQNSFRNQLEGFINGPQLHNRVHRWVGGQMGVVPTAPNDPVFFLHHANVDRIWAVWQIIHRNQNYQPMKNGPFGQNFRDPMYPWNTTPEDVMNHRKLGYVYDIELRKSKRSSLEHHHHHH | 34,521 |
| cGAS Enzyme (Sumo tagged) | **cGAS** | GSSHHHHHHSSGLVPRGSHMSDSEVNQEAKPEVKPEVKPETHINLKVSDGSSEIFFKIKKTTPLRRLMEAFAKRQGKEMDSLRFLYDGIRIQADQTPEDLDMEDNDIIEAHREQIGGMGASKLRAVLEKLKLSRDDISTAAGMVKGVVDHLLLRLKCDSAFRGVGLLNTGSYYEHVKISAPNEFDVMFKLEVPRIQLEEYSNTRAYYFVKFKRNPKENPLSQFLEGEILSASKMLSKFRKIIKEEINDIKDTDVIMKRKRGGSPAVTLLISEKISVDITLALESKSSWPASTQEGLRIQNWLSAKVRKQLRLKPFYLVPKHAKEGNGFQEETWRLSFSHIEKEILNNHGKSKTCCENKEEKCCRKDCLKLMKYLLEQLKERFKDKKHLDKFSSYHVKTAFFHVCTQNPQDSQWDRKDLGLCFDNCVTYFLQCLRTEKLENYFIPEFNLFSSNLIDKRSKEFLTKQIEYERNNEFPVFDEF | 55,741 |

***SI Figure S1.*** Peptide and protein sequences and properties. Molecular weights were computed using the Expasy Compute pI/Mw tool.

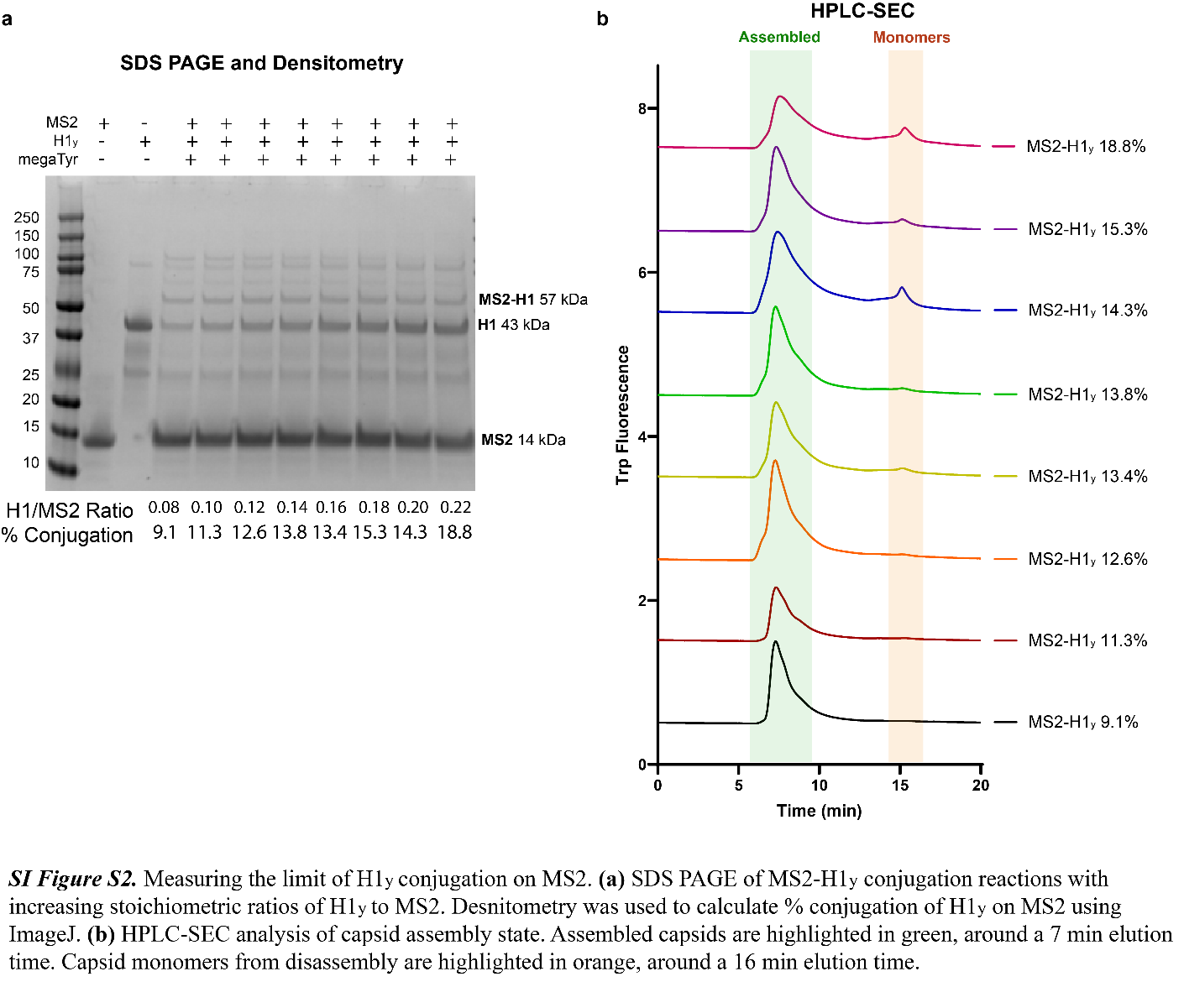

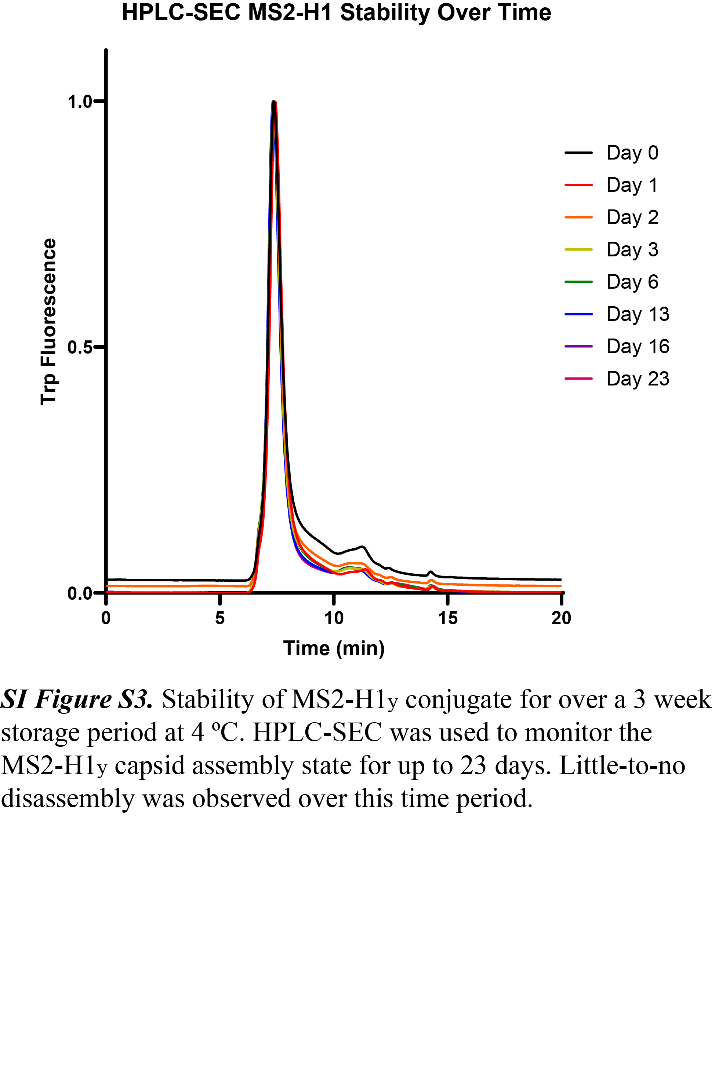

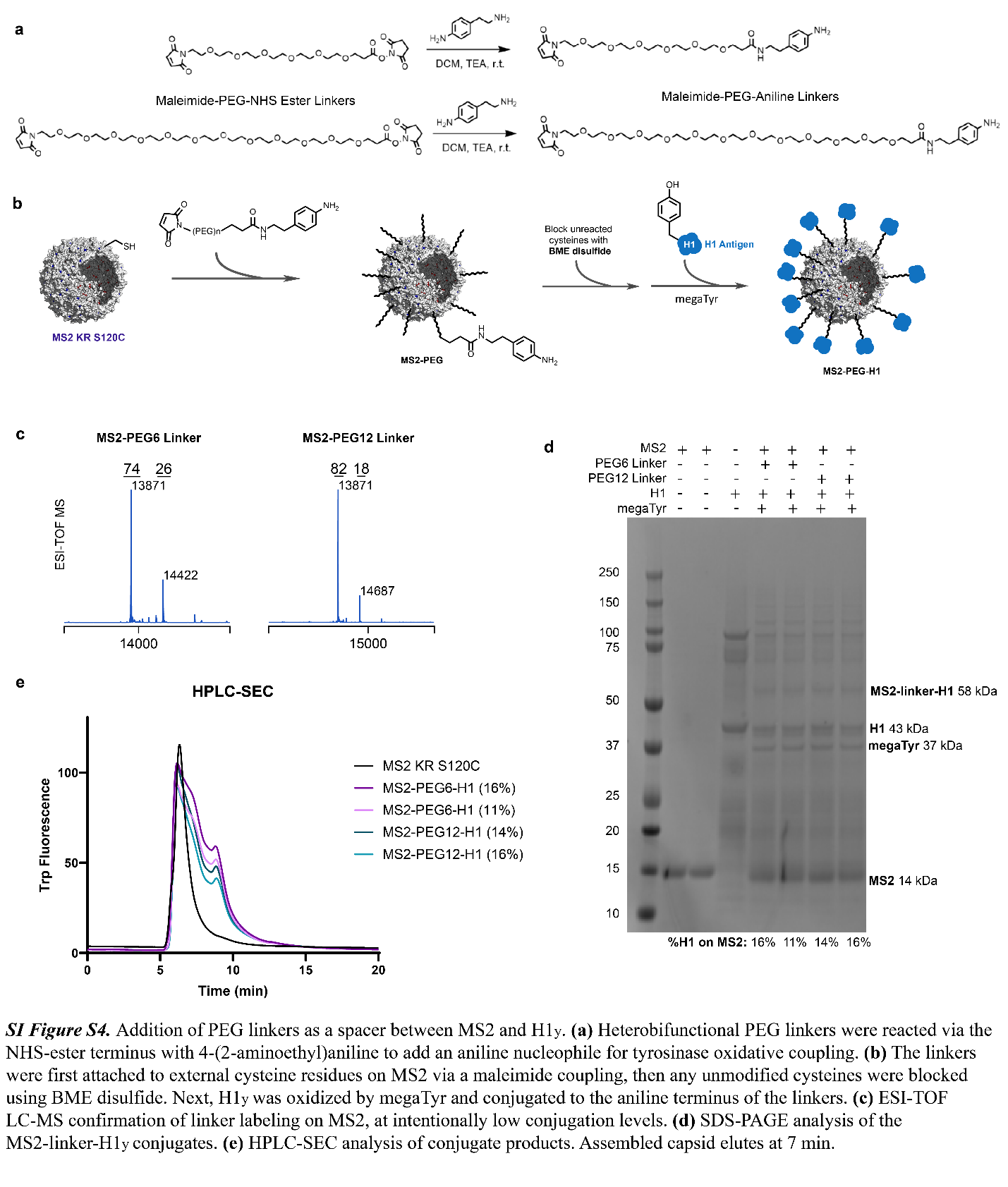

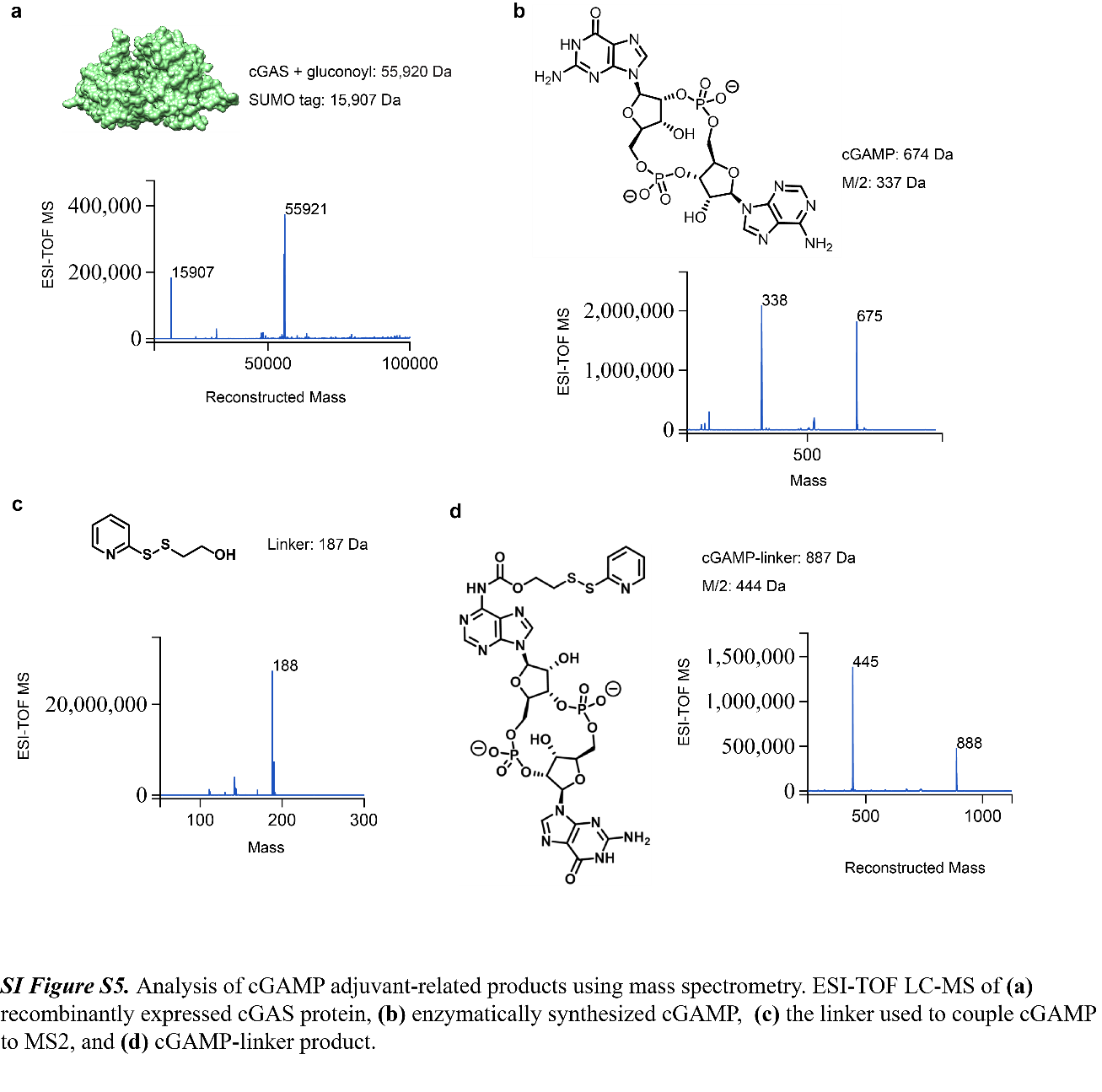

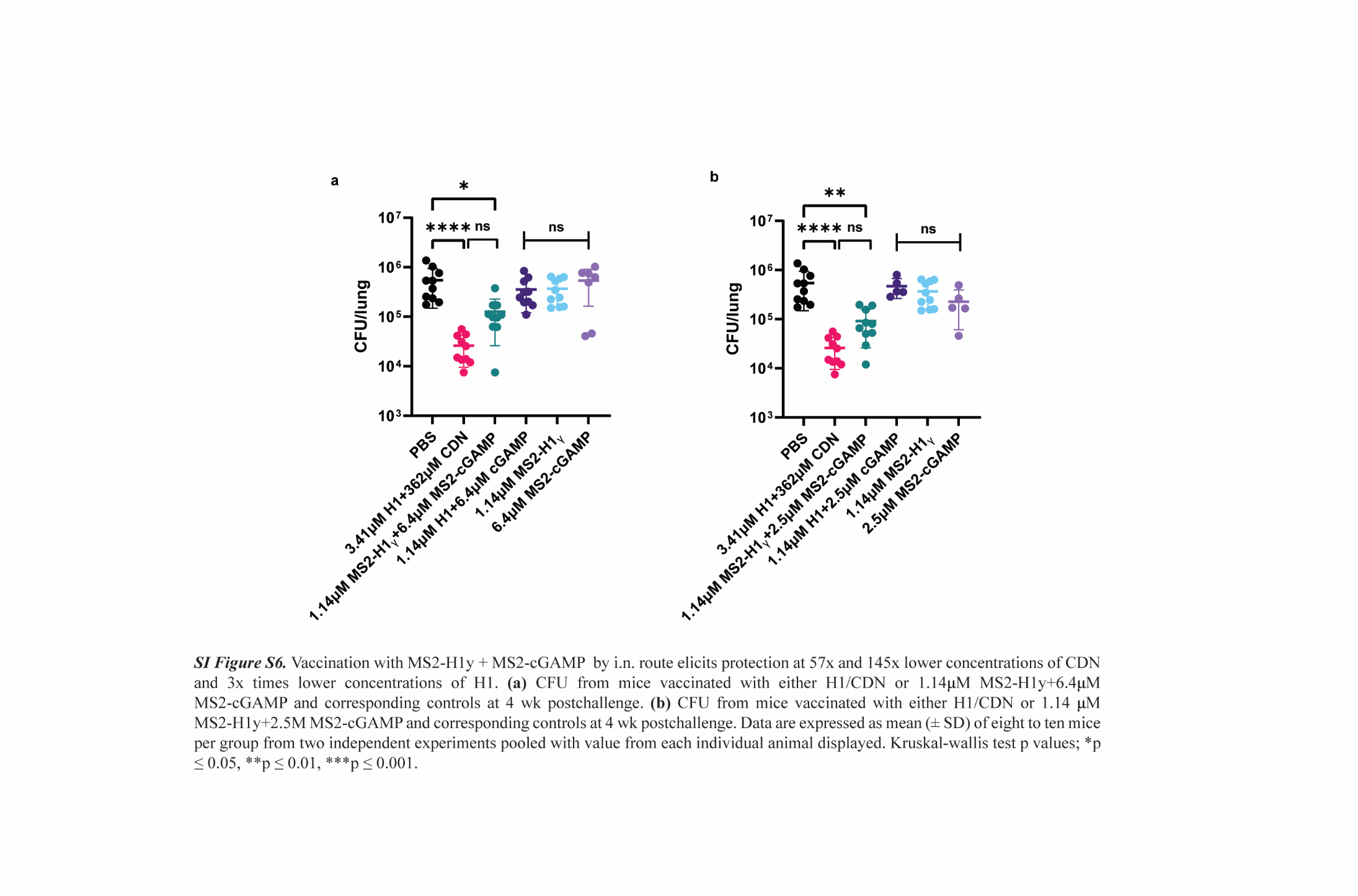
